# Common patterns of hydrolysis initiation in P-loop fold nucleoside triphosphatases

**DOI:** 10.1101/2022.06.23.497298

**Authors:** Maria I. Kozlova, Daria N. Shalaeva, Daria V. Dibrova, Armen Y Mulkidjanian

**Author notes:** **For correspondence:** Armen Y. Mulkidjanian, School of Physics, University of Osnabrueck, D-49069, Osnabrueck, Germany. **E-mail addresses:** Maria I Kozlova, Daria N. Shalaeva, Daria V. Dibrova, Armen Y. Mulkidjanian.

## Abstract

In ubiquitous P-loop fold <u>n</u>ucleoside <u>t</u>ri<u>p</u>hosphatases (also known as Walker NTPases), hydrolysis of ATP or GTP is initiated by interaction with an activating partner (usually another protein domain), which is accompanied by insertion of stimulatory moiety(ies) (usually arginine or lysine residues) into the catalytic site. After inspecting over 3600 Mg-NTP-containing structures of P-loop NTPases, we identified those with stimulator(s) inserted into catalytic sites and analysed the patterns of stimulatory interactions. In most cases, at least one stimulator twists gamma-phosphate counter-clockwise by linking the oxygen atoms of alpha- and gamma-phosphates; the twisted gamma-phosphate is stabilized by a hydrogen bond with the backbone amino group of the fourth residue of the Walker A motif. In the remaining cases, the stimulators only interact with gamma-phosphate. The ubiquitous mechanistic interaction of diverse stimulators with the gamma phosphate group suggests its twist/rotation as the trigger for NTP hydrolysis.

## 1. Introduction

Hydrolysis of nucleoside triphosphates (NTPs), such as ATP or GTP, by so-called P-loop fold nucleoside triphosphatases (also known as Walker ATPases) is one of the key enzymatic reactions. P-loop NTPases are coded by up to 20% gene products in a typical cell. P-loop NTPase domains drive the activity of rotary ATP synthases, DNA and RNA helicases, kinesins and myosins, ABC-transporters, as well as most GTPases, including ubiquitous translation factors, α-subunits of signaling heterotrimeric G-proteins and oncogenic Ras-like GTPases [1–15]. In the ECOD database [16], the topology-level entry “P-loop_NTPase” contains 193 protein families. In the Pfam database [17], the P-loop NTPase clan CL0023 contains 217 families. The main classes of P-loop NTPases, which are listed in Fig. S1 of Supplementary Figures and characterized in the accompanying article [18], were already present in the Last Universal Cellular Ancestor (LUCA) [7, 8, 13, 15, 19–23].

In P-loop NTPases, the eponymous P-loop, together with the first two residues of the following α-helix, usually follows the sequence pattern [G/A]xxxxGK[S/T], known as the Walker A motif [1]. This motif is stabilizing the triphosphate chain of NTP and the cofactor Mg^2+^ ion, see [1, 2, 24] and Fig. 1A-D. The Walker B motif hhhh[D/E], where “h” denotes a hydrophobic residue, is the other common motif of P-loop NTPases that is found on the C-terminal tip of that β-strand, which is opposite the Walker A α-helix, see Fig. 1A,B and [1, 2, 24]. The conserved Asp/Glu residue of the Walker B motif links it with the last [S/T] residue of the Walker A motif by a hydrogen bond (H-bond), see Fig. 1B-D.

**Figure 1.**
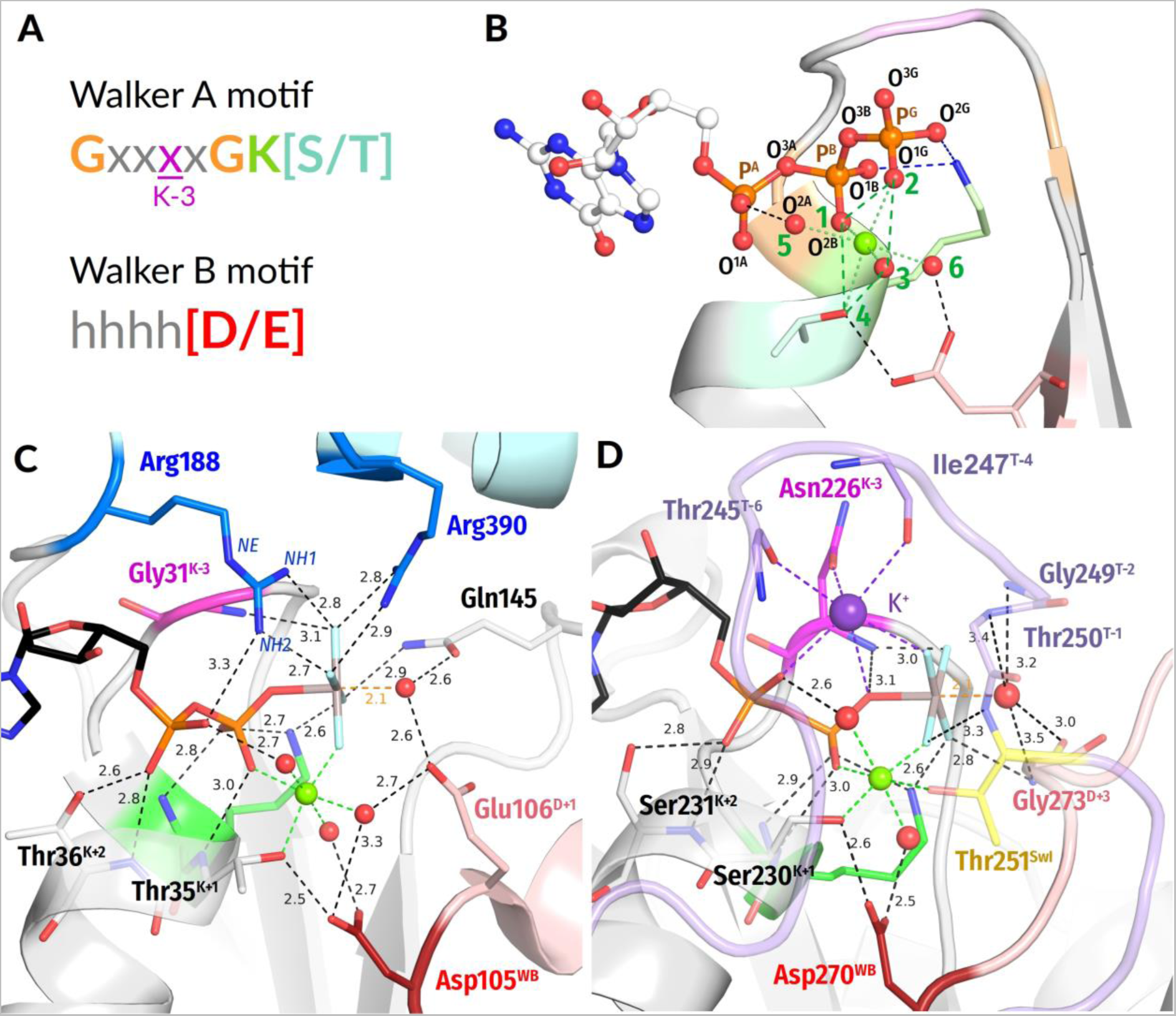
P-loop fold NTPases. **A**, conserved motifs in P-loop NTPases; **B**, naming of atoms according to IUPAC recommendations for nucleoside triphosphates [25] and typical Mg^2+^ coordination; **C, D,** crystal structures of typical catalytic sites of P-loop NTPases. **C,** Superfamily 1 helicase Pif1 with a transition state analog ADP-AlF_4_^-^ bound (PDB ID 5FHD, [26]). **D,** K^+^-dependent GTPase MnmE with a transition state analog GDP-AlF_4_^-^ bound (PDB ID 2GJ8 [27]). Color code: Polypeptide chains are shown as gray cartoons; nucleotides, their analogs, and important amino acid residues are shown as sticks, water molecules – as red spheres, Mg^2+^ ions are shown as lime spheres, K^+^ - as a purple sphere; the conserved Lys residue of Walker A motif is shown in green, and the amino acid residue three residues before it is highlighted in magenta; the conserved Asp residue of Walker B motif is shown in dark red, and the Arg fingers are shown in blue. In those amino acid residues that are shown as sticks, the oxygen atoms are colored red, and the nitrogen atoms are colored blue. In the AlF_4_^-^ moiety, the Al atom is colored gray, and the fluoride atoms are colored light blue. Switch I/K-loop motif is shown in lilac, the conserved Thr^SwI^ is highlighted in yellow.

An uncontrolled NTP hydrolysis would be perilous for cell survival. Therefore, a specific feature of most P-loop ATPases is their activation before each turnover. Usually, an ATP- or GTP-bound P-loop domain interacts with its cognate activating partner, which could be a domain of the same protein, a separate protein and/or a DNA/RNA molecule. Upon this interaction, specific stimulatory moieties, usually Arg or Lys residues (“Arg/Lys finger(s)”, Fig. 1C), are inserted into the catalytic site and initiate the cleavage of γ-phosphate [4, 12, 27–36]. As a rule, large-scale conformational changes accompany both the binding of NTP (“closing” of the catalytic pocket) and the NTP hydrolysis (“opening” of the catalytic pocket); these structural changes are coupled to useful mechanical work and drive most biomolecular motors, see, e.g. [4, 37–39]. In this paper, however, we focus only on the interaction of stimulators with the NTP molecules within catalytic pockets.

Important hints for clarifying the events inside these pockets are provided by their structures with bound transition state (TS) analogs, such as NDP:AlF_4_^-^ or NDP:MgF_3_^-^ or NDP-VO_4_^-^ complexes [4, 28, 29, 35, 40–44]. It is anticipated that these analogues promote the formation of a full-fledged active site in a pre-catalytic state. In such complexes, a water molecule is often located apically to the analog of γ-phosphate, which corroborates the earlier suggestions that γ-phosphate cleavage is triggered by the apical nucleophilic attack of a polarized water molecule (W_cat_), after its deprotonation to OH^—^_cat_, on γ-phosphate, see Fig. 1C, D; the accompanying article [18] and [12, 35, 44–49]. Also, TS analogs were shown to promote the interaction of the activator with the NTPase domain and therefore often disclose the interaction of the stimulator with the triphosphate chain, as shown in Fig. 1C-D, scrutinized in the accompanying article [18], and discussed in [28, 29, 35, 36, 40–42, 44, 50–59].

There is no consensus on how do stimulators initiate the hydrolysis. Warshel and colleagues suggested that stimulators, owing to their positive charge, compensate the negative charges of oxygen atoms of triphosphate and make the γ-phosphorus atom (P^G^) more prone to the nucleophilic attack by a water molecule [60–62]. It was also suggested that the pushing of bound water molecules out of the binding pocket into the bulk solution by the stimulatory arginine finger provides an entropic gain [63]. Jin and colleagues proposed reorientation of the W_cat_ molecule into the attack position by the stimulator [44, 64, 65]. On a later reaction step, stimulators were implied in compensating for the negative charge that develops at β-phosphate upon the breakaway of γ-phosphate [49, 66]. Recently, Gerwert and colleagues proposed, for small GTPases, that the stimulatory Arg finger rotates the α-phosphate group towards an eclipsed conformation with respect to β- and γ-phosphates, which would destabilize the triphosphate chain [67–69].

Elsewhere [70], we have focused on P-loop NTPases that are stimulated not by Arg/Lys residues but by K^+^ and Na^+^ ions, see [27, 33, 71–74]. We performed molecular dynamics (MD) simulations of the Mg^2+^-GTP-containing dimer of K^+^-dependent tRNA-modifying enzyme MnmE (see Fig. 1D) and compared their results with MD simulations of water-dissolved Mg^2+^-GTP and Mg^2+^-ATP complexes in the presence of monovalent cations. In the latter case, one of the monovalent cations bound between the O^2A^ and O^3G^ atoms of α- and γ-phosphates, in the so-called AG site, which was accompanied by rotation of α-phosphate - as the least constrained phosphate group - yielding a fully eclipsed configuration of the triphosphate chain [70]. These data agreed with the results of Gerwert and colleagues who used a Mg^2+^- metyltriphosphate complex – as a simple mimic of Mg^2+^-GTP – in their modeling the interaction with an Arg finger [67].

However, MD simulations of the GTP-containing MnmE dimer showed that α-phosphate was fully immobilized by bonds between the GTP molecule and the P-loop; these interactions stabilize the triphosphate chain in an extended, supposedly catalytically prone conformation see Fig. 1D and [70]. In this case, only the terminal γ-phosphate retained some mobility. The K^+^ ion, by binding between the O^2A^ and O^3G^ atoms of the protein-bound GTP molecule, caused twisting of the γ-phosphate group by approx. 30°; the rotation was counterclockwise when viewed from the side of γ-phosphate, see Fig. S2 and [70]. It is important that the K^+^ ion can hardly link these two oxygen atoms without twisting γ-phosphate. The twisted γ-phosphate was stabilized by a new H-bond between the O^2G^ atom of γ-phosphate and the backbone amide of Asn226, the fourth residue of the Walker A motif, see Fig. 1A, 1D, S2 and [70]. Both the twisting of γ-phosphate and its stabilization by an additional H-bond could potentially promote hydrolysis [70]. Still, these features were not discussed in relation to the catalytic mechanisms of any P-loop NTPase before. Therefore, we were interested to find out how widespread are the features we identified in K^+^-stimulated GTPase MnmE.

Here, we performed the first, to our best knowledge, comparative structure analysis of stimulation patterns in all relevant structures of P-loop NTPases. We screened 1484 available structures of P-loop NTPases with Mg-NTPs or their analogs bound in their catalytic sites (in total 3666 individual catalytic sites were analyzed). In this search, we relied on a key observation that diverse P-loop NTPases, despite playing dramatically different roles in the cell, catalyze essentially the same or similar chemical reactions [9]. It can be expected that catalysis of congruent chemical reactions in related enzymes is accomplished by homologous structural elements.

We have found two major types of interaction between stimulatory moieties and triphosphate chains. In most cases, as in the MnmE GTPase, at least one stimulator links O^2A^ and O^3G^, which enforces a counter-clockwise rotation of γ-phosphate. Otherwise, the stimulators interact only with the γ-phosphate group. In this case, the direction of twisting/pulling of γ-phosphate should be clarified on a case-to-case basis. Also, we noticed that not all stimulators carry a full positive charge; in diverse classes of P-loop NTPases we found asparagine, serine, and glycine residues forming stimulatory bonds with oxygen atoms of the triphosphate chain. In general, the only feature that seems to be common to all the studied P-loop NTPases is the mechanistic interaction of diverse stimulators with the γ-phosphate group.

This common feature allowed us to suggest that stimulators may initiate NTP hydrolysis by displacing the oxygen atoms of γ-phosphate, in particular, the O^1G^ atom that coordinates the Mg^2+^ ion. In the accompanying article [18] we further consider the functional consequences of the γ-phosphate twist and suggest a common catalytic mechanism for P-loop fold NTPases.

## 2. Results

### 2.1. Generic numbering of key amino acid residues for P-loop NTPases

Historically, different families of P-loop NTPases were studied by different research communities, each developing its own terminology. Therefore, no generic amino acid numbering for P-loop NTPases exists. For convenience, and by analogy with the Ballesteros-Weinstein numbering scheme for the G-protein coupled receptors [75], we introduce here a generic residue numbering for the conserved regions of P-loop NTPases. Without this generic nomenclature, our comparison of stimulatory mechanisms in diverse classes of P-loop NTPases, as performed here and in the accompanying paper of Kozlova et al. [18], would be hardly possible.

First, we chose highly conserved residues as benchmark references. In the Walker A motif, this is the conserved Lys residue (K^WA^) that forms H-bonds with the O^1B^ oxygen atom of β-phosphate and O^2G^ atom of γ-phosphate, see Fig. 1. In the Walker B motif, we have chosen the conserved [Asp/Glu]^WB^ residue ([D/E]^WB^) that makes an H-bond with the last Ser/Thr residue of the Walker A motif (see Fig. 1) and is involved in the coordination of the Mg^2+^ ion.

In addition to universal Walker A and Walker B motifs, we also invoked the Switch I motif that is located between the Walker A and Walker B motifs in NTPases of TRAFAC (TRAnslation FACtors) class, see Fig. S1 and [7, 76]. The Switch I has only a single strictly conserved [Thr/Ser]^SwI^ residue ([T/S] ^SwI^, colored yellow in Fig. 1D) that can be used as a reference. In NTPases of the TRAFAC class, the side chain of this [T/S]^SwI^ residue coordinates the Mg^2+^ ion, its backbone amino group (hereafter **HN**) forms a H-bond with γ-phosphate, and its backbone carbonyl group (hereafter **CO**) stabilizes W_cat_, see Fig. 1D.

In the following, we number the amino acids of the Walker A, Switch I, and Walker B motifs relatively to the reference residues, as shown in Fig. 1. In this case, the Asp226 residue of MnmE GTPase, the fourth residue of Walker A motif, which binds the K^+^ ion by its side chain and makes a H-bond with O^2G^ via its backbone **HN** group (see Fig. 1D, S2 and [70]), is denoted as Asp226^K-3^ or Asp^K-3^. Its backbone **HN** group proper is then labeled as **HN**^K-3^.

To distinguish the interaction of the P-loop domain with its cognate activating partner from the consequent insertion of, say, an Arg finger into the catalytic site, we consistently refer to those elements that are poked into catalytic sites to stimulate hydrolysis as ’stimulatory moieties’ or ’stimulators’; see, for instance, the dark blue Arg fingers in Fig. 1C or the K^+^ ion in Fig. 1D. We follow Wittinghofer and colleagues who wrote about “GAP stimulated GTPase activity” in one of their pioneering works [77]. Furthermore, the original Latin meaning of “stimulus” – “a sharp stick used to poke cattle to get them to keep moving” (quoted from https://www.dictionary.com/browse/stimulus) - nicely describes the function of Lys and Arg fingers in P-loop NTPases that routinely empower movements of cellular structures.

### 2.2. Global structural analysis of stimulatory patterns in the whole set of P-loop NTPases with bound Mg-NTP complexes or their analogs

The ever-rising four-digit numbers of P-loop NTPase structures with ATP, GTP, and their analogs bound, as deposited in the PDB, demanded computational approaches. The search in the Protein Data Bank (PDB) at www.rcsb.org [78, 79] for proteins assigned to the entry IPR027417 “P-loop containing nucleoside triphosphate hydrolase” of the Interpro database [80] yielded as many as 1484 structure entries with 3666 catalytic sites with NTP or NTP-mimicking molecules bound (as of 11.09.2019; many of the structures contained several catalytic sites). The criteria for selection of full-fledged catalytic sites from this set and the routine of their subsequent structural analysis are described and depicted in the Methods section. After filtering, we obtained 3136 structures of catalytic sites containing complexes of Mg^2+^ ions with NTPs or NTP-like molecules, these structures were subjected to further analysis. The relevant data for all these catalytic sites are presented in Supplementary Tables 1 and 2 (hereafter Tables S1 and S2, data as of 11.09.2019).

Based on the type of the molecule, the complexes could be sorted into four groups: 1043 sites contained native ATP/GTP molecules; 1612 sites contained bound non-hydrolyzable NTP analogs such as Adenosine 5′-[β,γ-imido]triphosphate, (AMP-PNP), Guanosine 5′-[β,γ-imido]triphosphate (GMP-PNP), Adenosine 5′-[*β*,*γ*-methylene]triphosphate (AMP-PCP), Guanosine 5′-[*β*,*γ*-methylene]triphosphate (GMP-PCP), Adenosine 5′-[γ-thio]triphosphate (ATP-γ-S), and Guanosine 5′-[γ-thio]triphosphate (GTP-γ-S); 234 sites contained NDP:fluoride complexes mimicking the substrate state, such as NDP:BeF_3_ and NDP:AlF_3_, and 247 sites contained NDP:AlF_4_^-^(204), NDP:MgF_3_^-^ (10) and ADP:VO_4_^-^ (33) thought to be TS analogs [35, 40, 42–44].

#### 2.2.1. Stabilization of the O^2G^ atom of γ-phosphate by HN^K-3^ of the Walker A motif

For the MnmE GTPase, we have shown that the insertion of a K^+^ ion and its simultaneous interaction with O^2A^, O^3B^, and O^3G^ atoms triggered the twist of γ-phosphate, leading to the formation of a new H-bond between the O^2G^ atom and **HN** of Asn226^K-3^, see Fig. 1D, S2 and [70]. Generally, the position of **HN**^K-3^ in the vicinity of the O^2G^ atom (or the corresponding atom of an NTP analog) is structurally conserved across P-loop NTPases, being determined by the highly conserved H-bond of **HN**^K-3^ with the bridging O^3B^ oxygen (see Fig. 1C,D).

To assess the possibility of a transient H-bond formation between **HN**^K-3^ and O^2G^ of γ-phosphate, as found in MD simulations of MnmE GTPases [70], we measured corresponding distances in the available structures of P-loop NTPases with bound substrates or their analogs as described and depicted in Methods. The data obtained are presented in Table S1 and Fig. 2, where the H-bond-compatible distance range is highlighted in amber. For simplicity, we used the same threshold of 3.4 Å for the H-F and H-O bonds. On the one hand, this value is somewhat lower than the threshold of 3.5 Å, as suggested for H-bonds in protein structures by Martz [81]. On the other hand, this distance corresponds to the longest F-H-N bond reported for crystalized L-cysteine-hydrogen fluoride [82]. Still, this threshold is rather arbitrary; according to Jeffrey, weak H-bonds in proteins can have donor-acceptor distances up to 4.0 Å long [83].

**Figure 2.**
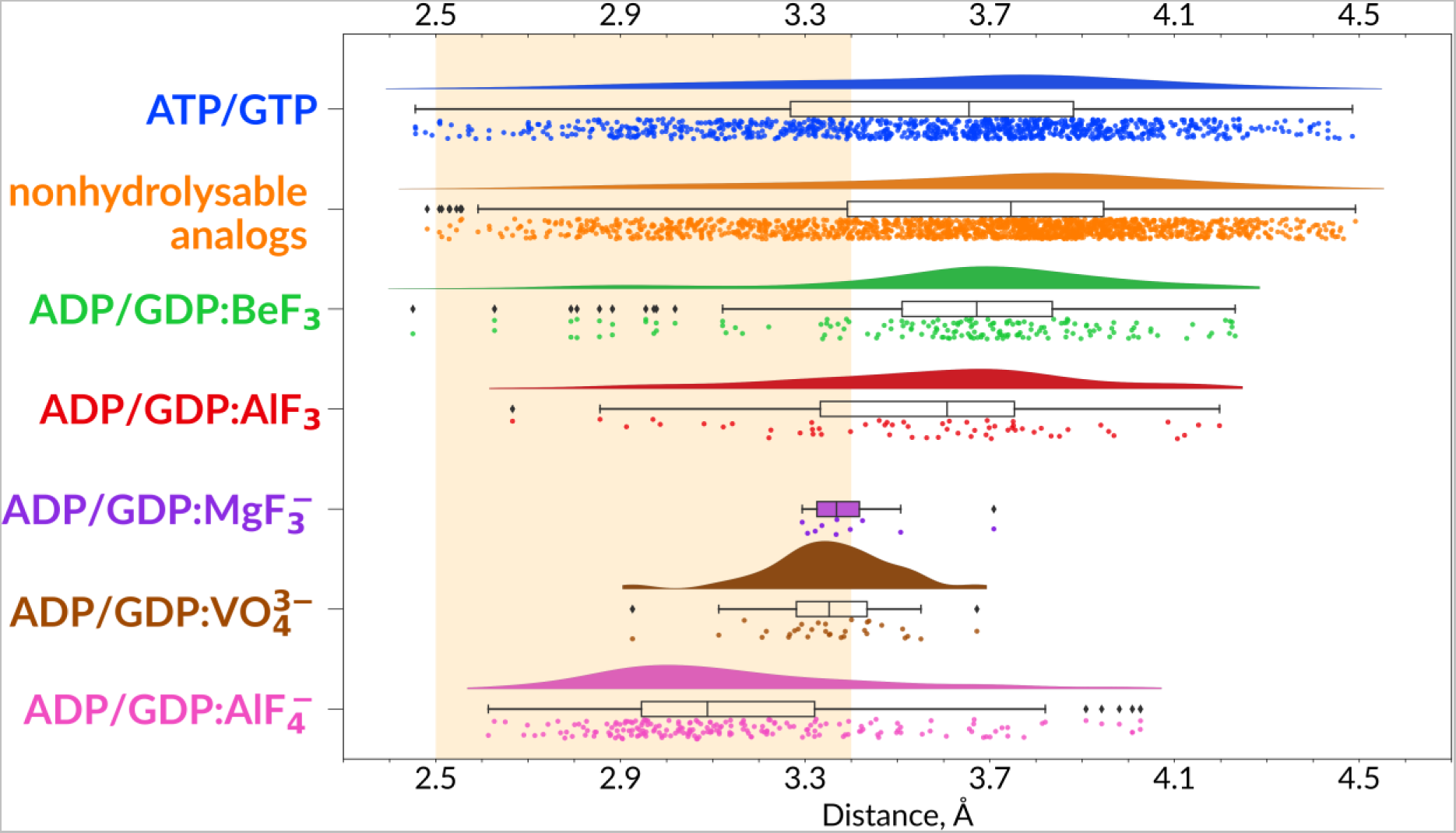
Distances between HN^K-3^ and the O^2G^ atom or its analog in the catalytic sites of P-loop fold NTPases. For each type of complexes, distances are visualized as a kernel density estimate (KDE) plot, a boxplot, and individual data points, each point representing one catalytic site in one structure. For ADP/GDP:MgF_3_^-^ complexes the density plot is not shown because of scarcity of data. The range of H-bond-compatible lengths is highlighted in amber.

In all groups of complexes, there is a fraction with distances shorter than 3.4 Å between **HN**^K-3^ and the nearest O^G^ atom or its structural analog (hereafter **HN**^K-3^ - O^G^ distance). For the ATP- and GTP-containing structures, this fraction makes 31% of complexes (326 of 1043 binding sites); for the non-hydrolyzable analogs, it makes 24% of complexes (392 out of 1612 binding sites). Among complexes with NDP:BeF_3_, the **HN**^K-3^ - O^G^ distance is < 3.4 Å in 20.5% of cases(35 catalytic sites out of 171), whereas in the case of structures containing NDP:AlF_3_ this fraction makes 28.5% of catalytic sites.

However, the distances between **HN**^K-3^ and O^G^ analog are shorter than 3.4 Å in most structures with TS analogs as shown in Fig. 2 where the data for the three TS analogs are plotted separately for clarity. Distances of < 3.4 Å are observed in 61% of complexes with ADP:VO_4_^3-^ (out of 33), in seven out of ten NDP:MgF_3_^-^ complexes and in 80% of NDP:AlF_4_^-^ complexes (163 out of 204). Hence, the TS-like structures of catalytic sites correlate with a H-bond compatible distance between **HN**^K-3^ and O^G^ analog.

#### 2.2.2. Pre-catalytic configurations in NTP-containing structures

While H-bond-compatible distances between **HN**^K-3^ and O^G^ (or its analog in the case of TS- like structures, Fig. 2) confirmed our earlier suggestion on the importance of this H-bond for stabilization of the TS [70], we were rather surprised to see that the **HN**^K-3^–O^G^ distances were H-bond compatible also in 31% of ATP- and GTP-containing structures (Fig. 2, Table S1). Therefore, we inspected the top 100 high-resolution NTP-containing structures with H-bond compatible **HN**^K-3^–O^2G^ distances to clarify their origin.

In principle, an ATP or GTP molecule could be crystalized within an NTPase only if the latter is inactive. Not surprisingly, the ATP or GTP-containing P-loop NTPases which were crystalized in the absence of their cognate activators made the majority in the inspected set. Although in these proteins the **HN**^K-3^–O^2G^ distances were indeed H-bond compatible, we did not explore these structures further because they could hardly help in clarifying the mechanisms of hydrolysis stimulation.

Still, we could identify a set of structures where NTP molecules remained not hydrolyzed despite the presence of activating partner(s). In many such complexes, the interaction of W_cat_ with γ-phosphate was hindered by the incompleteness of W_cat_ ligands, specifically caused by mutations, so that the NTP-binding sites were trapped in precatalytic configurations. Some such structures are shown in Fig. 3A-C. In these enzymes, the stimulatory fingers are inserted into the catalytic sites, γ-phosphates are twisted, and the triphosphate chains are in a configuration like that observed upon the MD simulations of MnmE GTPase [70], which is superimposed as a dark-red contour (cf Fig. S2). One can see from Fig. 3A-C that H-bond compatible **HN**^K-3^-O^2G^ distances correlate with linking of O^2A^ and O^3G^ atoms by the stimulator, twist of γ-phosphate, and a more eclipsed configuration of the triphosphate chain.

**Figure 3.**
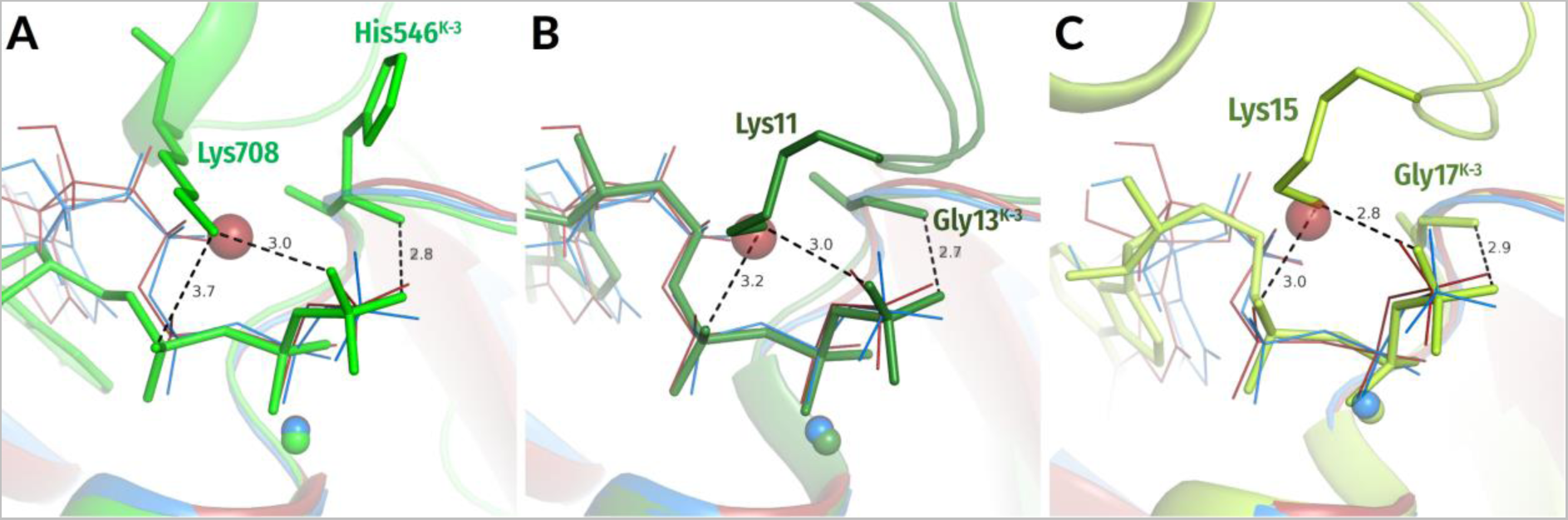
Conformation of native ATP molecules in selected crystal structures as compared to MD simulations of the MnmE GTPase. Crystal structures of P-loop ATPases in complex with ATP molecules are shown in lime or green colors; they are superposed with conformations of GTP bound to the MnmE GTPase, as sampled from MD simulations of inactive monomer (blue) and active, K^+^-bound dimer (red), see also Fig. S2 and [70]. All distances are given in ångströms. **A**. N-ethylmaleimide-sensitive factor (PDB ID 1NSF, [84]). **B**. ATPase MinD (PDB ID 3Q9L, [85]). **C**. Chromosome segregation protein Soj (PDB ID 2BEK, [86]).

#### 2.2.3. Geometry of the ADP:AlF_3_ complex in a P-loop NTPase

It is noteworthy that a particularly flat distribution of distances between NH^K-3^ and O^G^ analog is observed with the AlF_3_-containing complexes (Fig. 2). This flattness could be due to the suggested presence of NDP:MgF_3_^-^ complexes in some of the structures deposited in the PDB as NDP:AlF_3_-containing structures [35, 44, 58]. Earlier, it was shown that, depending on pH, aluminum fluorides can make complexes with NDP in two forms, yielding NDP:AlF_4_^-^ or NDP:AlF_3_ complexes [54]. However, after identification of NDP:MgF_3_^-^ as one more TS analog [42], Blackburn and colleagues argued that all NDP:AlF_3_ complexes from previously determined structures are, in fact, misassigned NDP:MgF_3_^-^ complexes [35, 44, 58]. They proposed that the low Al(OH)_3_ solubility above pH 7.5 would trigger the substitution of Al^3+^ for Mg^2+^ that is usually present in the crystallization solution of NTPases in high amounts to promote the NTP binding [35, 44]. Since the atomic numbers of Al and Mg atoms are very similar, specific methods, such as proton-induced X-ray emission spectroscopy (PIXE), are required to determine whether the structure contains NDP:AlF_3_ or a TS-analog NDP:MgF_3_^-^. Application of these methods to the crystals that were studied many years ago is hardly possible.

We manually inspected all the moieties denoted as AlF_3_ in the sampled structures of P-loop NTPases (Table S1). Upon the inspection, we identified an AlF_3_:ADP-containing structure of the Zika Virus helicase with a resolution of 2Å, see Fig. 4A and PDB ID 5Y6M, [87]. This structure could not contain a MgF_3_^-^ ion instead of AlF_3_ because the crystallization solution contained no Mg^2+^, but only Mn^2+^ ions. No manganese fluoride complexes with more than two F^-^ ions in the Mn^2+^ coordination sphere could be found in the PDB (as of 25.06.2020). Also, the inspection of electron density (Fig. 4B) ruled out the possibility of a misassigned AlF_4_^-^.

**Figure 4.**
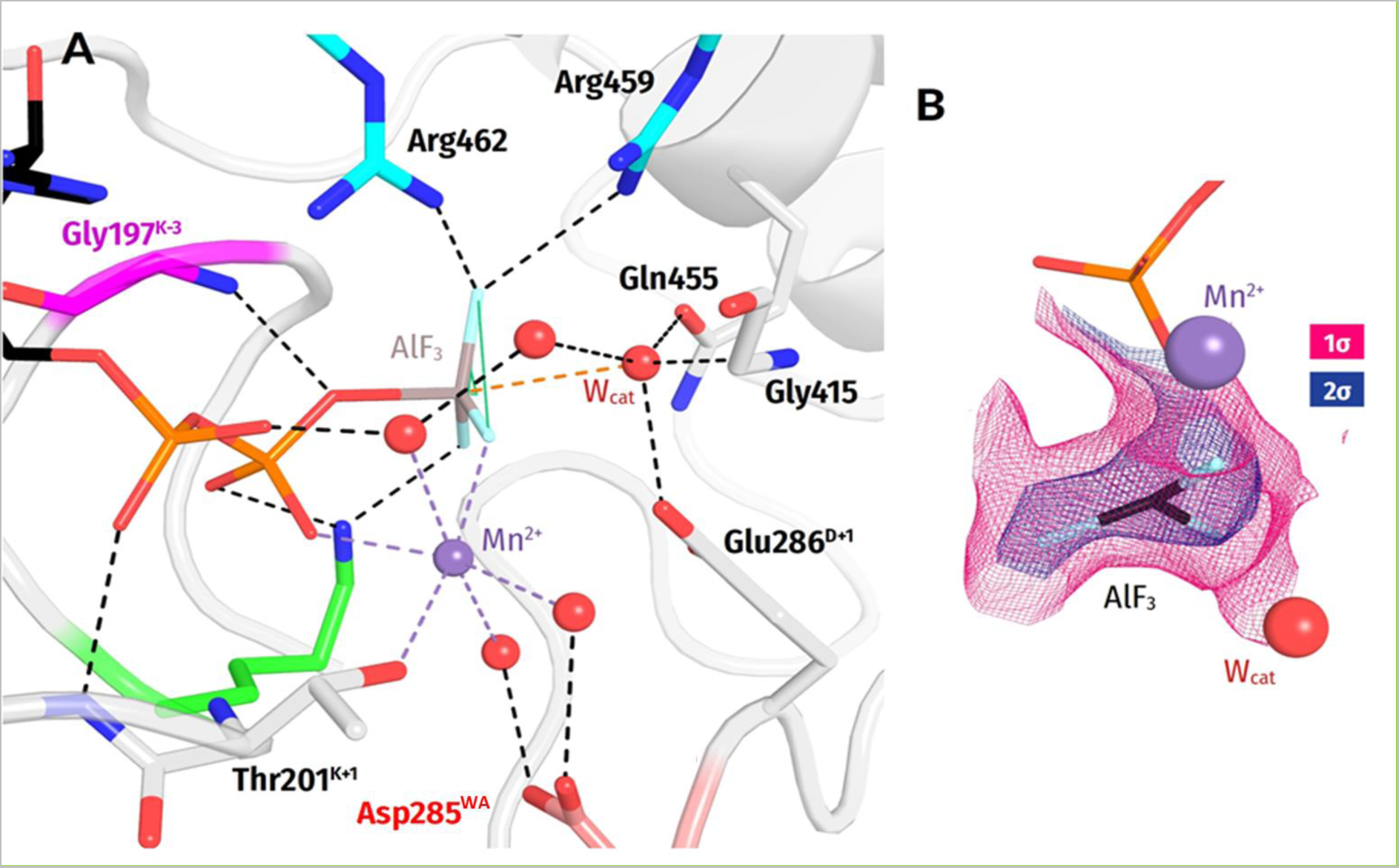
Zika virus helicase in complex with an ADP:AlF_3_ moiety. **A**. Nucleotide-binding pocket of Zika virus helicase with a ADP:AlF_3_ moiety bound (PDB ID 5Y6M [2]). Water molecules are shown as spheres. Mn^2+^ ion is shown as a purple sphere; the conserved Lys200^WA^ is shown in green, the Gly197 k-and the amino acid residue three residues before it is highlighted in magenta. The conserved Asp residue of Walker B motif is shown in pale red; the Arg fingers are shown in cyan. **B**. 2fo-fc electron density for AlF_3_ moiety contoured at 1σ and 2σ.

The high resolution of this helicase structure allowed us to determine the geometry of an Mn^2+^:ADP:AlF_3_ complex in a P-loop ATPase (Fig. 4). The F^3^-F^1^-F^2^-Al dihedral angle is 17.5° and the distance between the Al atom and W_cat_ is almost 3 Å, as compared with about 2.5 Å in MgF_3_^-^-containing complexes and about 2.0 Å in AlF_4_^-^-containing complexes [35, 44]. The **HN**^K-3^-O^2G^ distance was 3.97Å, i.e. longer than, on average, in complexes with MgF_3_^-^ and AlF_4_^-^ (cf. Fig. 2). This difference is unlikely to be due to the replacement of Mg^2+^ by Mn^2+^ since the radii of the two ions are similar.

Hence, our analysis indicates that P-loop-bound NDP:AlF_3_ complexes are present in the PDB. Their AlF_3_ moiety appear to have a substrate-like geometry that differs from that of planar, TS-like MgF_3_^-^ moieties. Hence, the mixing of NDP:AlF_3_ and NDP:MgF_3_^-^ complexes might indeed contribute to the flatness of distribution of distances to **HN**^K-3^ in the case of AlF_3_-containing structures in Fig. 2.

#### 2.2.4. Different modes of AlF_4_^-^ interaction with the Mg^2+^ ion

Also we noticed the breadth of distance distributions as measured in NDP:AlF_4_^—^-containing structures (Fig. 2). Of course, the nucleotide-binding pockets of diverse of P-loop NTPases could differ somewhat; the main classes of these enzymes, as shown in Fig. S1, split even before the LUCA [7, 8, 13, 15, 19–23, 88]. Also, the possible artifacts of structure determination and uncertainties of such determination (as specifically addressed in the Supplementary File 1) should be accounted for. Still, the manual examination of the NDP:AlF_4_^-^-containing structures revealed one more reason for the structural differences. It turned out that AlF_4_^-^ moieties can interact with Mg^2+^ in two different ways, at least, see Fig. 5A-D and Supplementary Table 3 (hereafter Table S3). In most structures (77% of all AlF_4_^-^ complexes), only one fluorine atom interacts with Mg^2+^, like its structural counterpart, the O^1G^ atom of γ-phosphate. In this case, the next closest fluorine atom is on > 3.0 Å from Mg^2+^, see Fig. 5A, 5C and Table S3. In some structures, however, the two fluoride atoms are at similar distances of 2.0-2.7 Å from the Mg^2+^ ion and both appear to interact with it, which is only possible when the AlF_4_^-^ is rotated by approx. 45° around the O^3B^-Al bond, see Fig. 5B, 5D and Table S3. This interaction is non-physiological because it usually prevents Mg^2+^ from interaction with one of its physiological ligands (see Fig. 5B and Table S3). Our manual inspection of the AlF_4_^-^ structures showed that, in general, structures with a second fluorine atom found within 3 Å from Mg^2+^ (these include structures with different degrees of AlF_4_^-^ rotation) usually have distortions in the Mg^2+^ coordination sphere, namely missing ligands, incorrect bonding to β-phosphate or a missing bond to Ser^K+1^. Only two catalytic sites out of 14 where two F—Mg distances are shorter than 2.7 Å appear not to have additional distortions (see Table S3). One of these two structures has a large Ca^2+^ ion instead of Mg^2+^ so that interactions in the coordination sphere are preserved due to a larger ionic radius of Ca^2+^ and its ability to bind up to 8 ligands (Fig. 5D).

**Figure 5.**
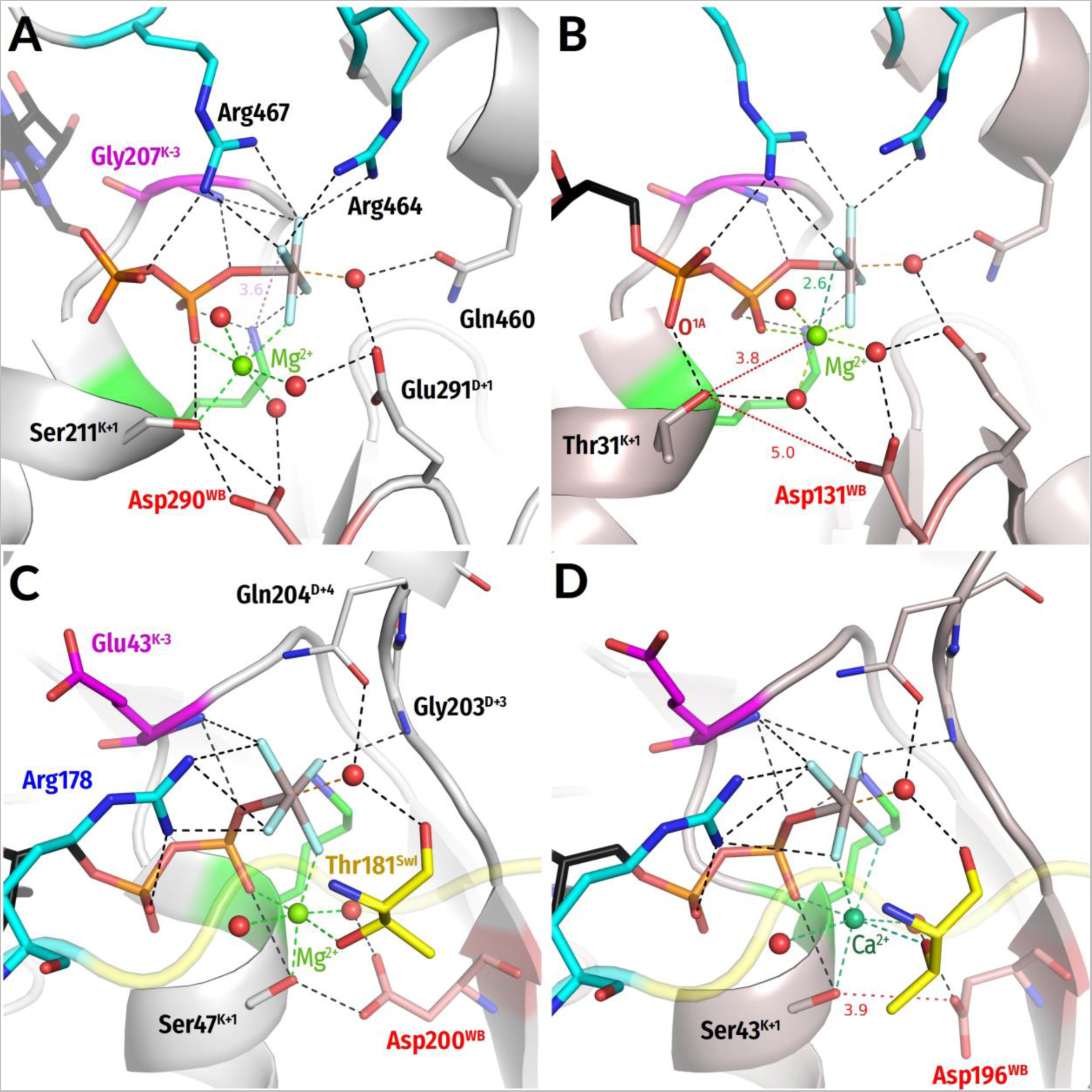
Variation in the AlF_4_^-^ interaction with Mg^2+^ ions. Residues are highlighted as in Fig. 1C,D. The Switch I motif is colored yellow, Mg^2+^ ions are shown as bright green spheres, Ca^2+^ ion is shown as a teal-colored sphere. The distances that are too long for H-bonds are indicated by red dashed lines. All distances are given in ångströms. **A.** The ADP:AlF_4_^-^ binding in the structure of HCV (Hepatitis C Virus) NS3 helicase, PDB 5E4F, chain A [89]. The Mg^2+^ ion is coordinated by the O^2B^ atom of ADP, one fluorine atom of AlF_4_^-^, the conserved Ser residue of Walker A motif, and three water molecules. **B.** The ADP:AlF_4_^-^ binding in the structure of double-stranded RNA (dsRNA)-dependent helicase LGP2 from chicken (PDB 5JAJ [90]). Here, AlF_4_^-^ is rotated counterclockwise and makes two bonds with the Mg^2+^ ion; the Ser^K+1^ residue is no longer directly coordinating Mg^2+^ and, instead, is forming an unusual bond with the O^1A^ atom of ADP. **C.** The AlF_4_^-^ binding in a structure of guanine nucleotide-binding protein Gα_i3_ complexed with the regulator of G-protein signaling 8 (PDB 2ODE [91]). The Mg^2+^ ion is coordinated by the O^2B^ atom of GDP, one fluorine atom of AlF_4_^-^, the conserved Ser residue of Walker A motif, the conserved Thr of Switch I, and two water molecules. **D.** In the structure of transducin Gtα complexed with Ca^2+^ instead of Mg^2+^ (PDB 1TAD [29]), AlF_4_^-^ is rotated and forms two bonds with Ca^2+^, but all interactions in the coordination sphere are preserved due to a larger ionic radius of Ca^2+^ and its ability to bind up to 8 ligands. However, the catalytic site is more open, as evidenced by a distance of 3.9 Å between Ser^K+1^ and Asp^WB^.

Accordingly, all structures with two bonds between fluoride atoms and Mg^2+^ (indicated in Table S3) appear to be suspicious as TS-state analogs because of non-physiological coordination of the Mg^2+^ ion; this point is addressed in the Discussion section and also the accompanying article [18].

Still, the comparison of the same or closely related NTPases with differently bound NDP:AlF_4_^-^ complexes has proven useful. Fig. 5A, 5B shows two structures of the Family 2 helicases (ASCE division, SF1/SF2 class, see Fig. S1) with different coordination of the AlF_4_^-^ moiety. One can see that the non-physiological coordination in Fig. 5B is achieved via counterclockwise rotation of the more “physiological” configuration of the AlF_4_^-^ moiety in Fig. 5A, whereby the interactions of AlF_4_^-^ with Lys^WA^ and the stimulatory Arg residues are retained in both structures. The two stimulatory fingers retain their H-bonds with AlF_4_^-^ in both configurations. This comparison shows that the residues which bind γ-phosphate appear to be adapted to the counterclockwise rotation of γ-phosphate by 30-40°. Similar counterclockwise rotation of γ-phosphate can be inferred from the comparison of G-α protein structures (Kinase-GTPase division, TRAFAC class, see Fig. S1) in Fig. 5C-D.

The data on all found non-physiologically bound AlF_4_^-^ moieties are highlighted pink in Table S1 and separately summarized in Table S3.

In sum, the data in Fig. 2, 3 and 5 indicate that the insertion of a stimulator and linking the O^2A^ and O^3G^ atoms leads to the twist of γ-phosphate and shortening of the **HN**^K-3^-O^2G^ distance in diverse NTP-containing P-loop NTPases in support of our earlier MD simulation data [70].

#### 2.2.5. Patterns of interaction between stimulatory moieties and the phosphate chain

In most families of P-loop NTPases, Arg, Lys, or Asn residues serve as stimulatory moieties. We used computational approach to inspect the patterns of their interactions with phosphate chain atoms (or their analogs) in the PDB-deposited structures of P-loop NTPases.

For this purpose, we analyzed, as described in the Methods section, the same 3136 PDB structures of catalytic sites that contain complexes of Mg^2+^ ions with NTPs or NTP-like molecules. For each complex, we measured the distances between oxygen atoms of the triphosphate chain (or their structural counterparts in the NTP analogs) and the amino groups of Arg, Lys, and Asn side chains within 4 Å radius, see the Methods section for further details and the scheme of the analysis pipeline. The distances were measured towards the NE/NH1/NH2 atoms of Arg residues; the NZ atom of Lys residues; and the ND2 atom of Asn residues (see Fig. 6 and 7 for the atom naming scheme). The data on all atom pairs and corresponding distances are summarized in Table S1.

**Figure 6.**
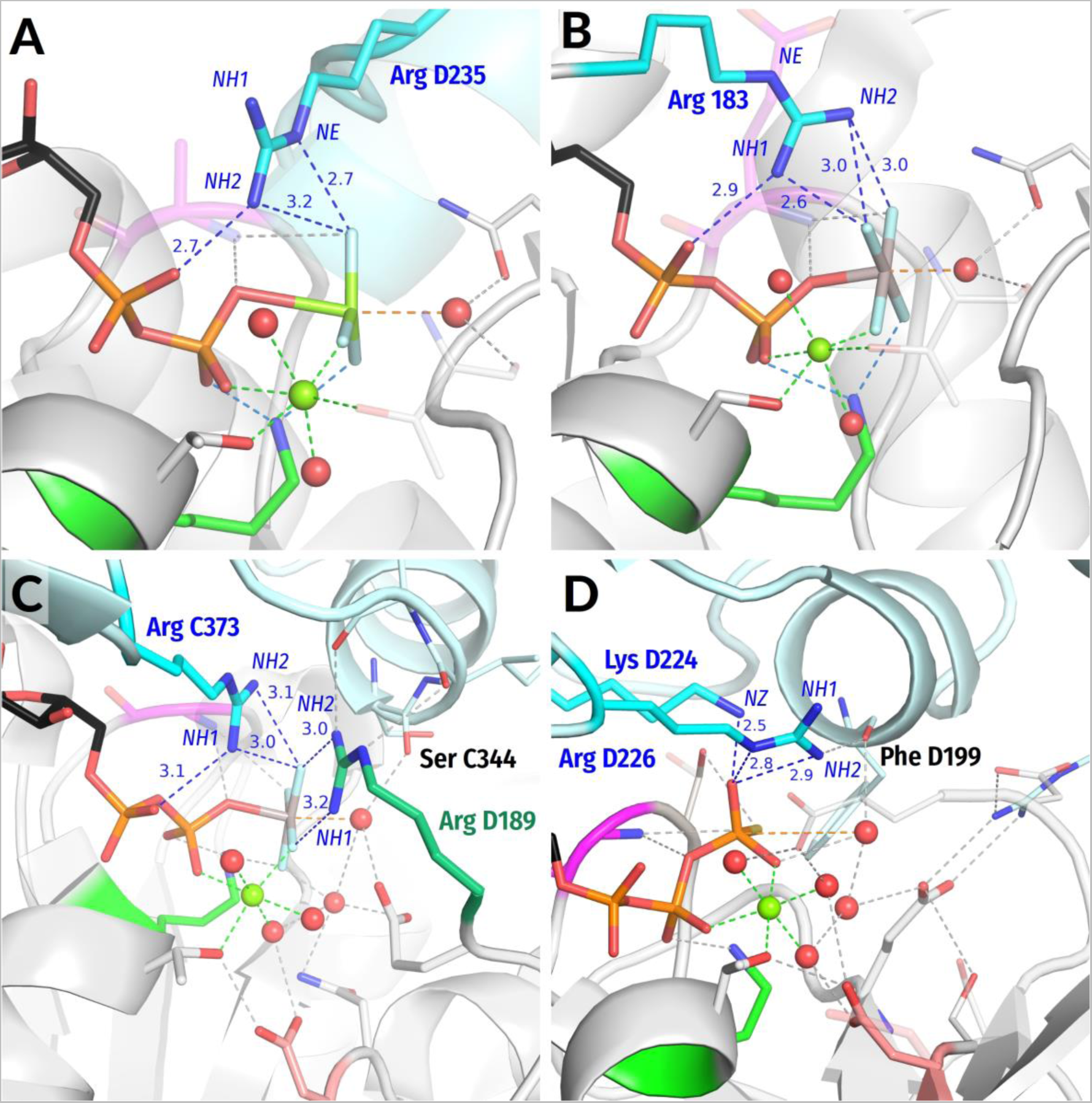
Examples of different interaction types/stimulatory patterns involving Arg residues. Protein fragments including P-loop domain are shown as grey cartoons, other subunits carrying Arg/Lys fingers are shown in cyan; functionally relevant residues are shown as sticks: P-loop Lys residue is shown in green, Arg and Lys fingers are shown in cyan; Mg cations are shown as green spheres. All distances are given in ångströms. **A**. Both α and γ phosphates are coordinated by the NH2 atom, an additional H-bond is formed by NE atom (stimulatory pattern AG); the structure of the Ras-like GTPase RhoA (PDB ID 5HPY, chain B [94]). **B**. Both α and γ phosphates are coordinated by NH1 atom, an additional H-bond is formed by the NH2 atom (stimulatory pattern AG); the structure of GTP-binding protein G(q) subunit α, (PDB ID 5DO9, chain A [95]). **C.** Both α and γ phosphates are coordinated by the NH1 atom, an additional H-bond is formed by NH2 atom (stimulatory pattern AG). An additional Arg residue provides more interactions with γ phosphate. Notably, this residue also interacts with the backbone atom of the α-subunit that provides the main Arg finger; the structure of bovine mitochondrial F_1_-ATPase, β-subunit (PDB 1H8E, chain D [41]). **D.** Only the γ-phosphate is coordinated by the NH2 atom of the Arg finger, the NE atom and Lys residue provide additional H-bonds (stimulatory pattern G_multi_); the structure of circadian clock protein KaiC, (PDB 4TL8, chain C [96]).

**Figure 7.**
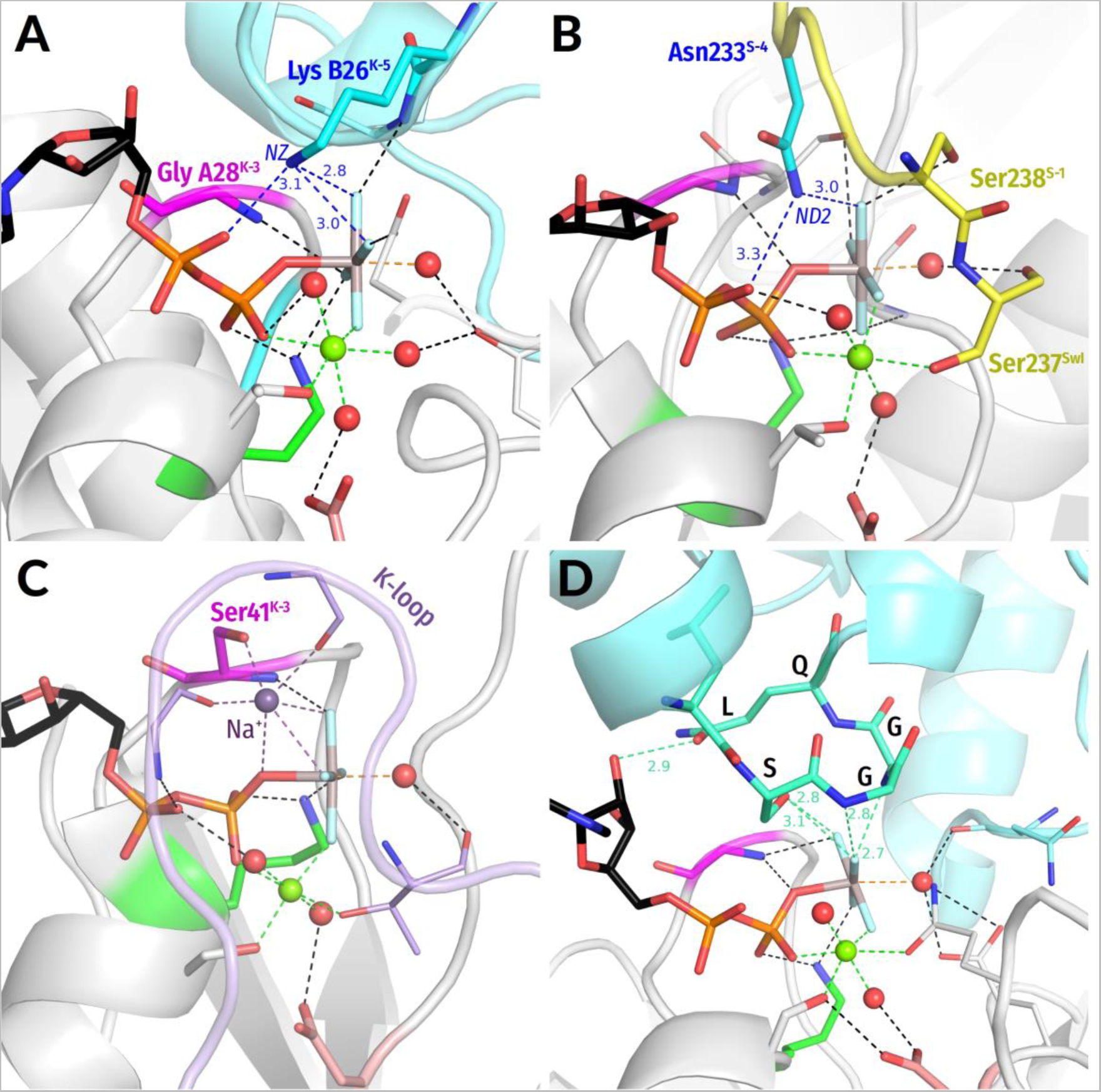
Examples of stimulatory interactions involving moieties other than arginine residues as stimulators. Residues are highlighted as in Fig. 6. **A**. Interaction of a Lys finger, as provided by another subunit of the homodimer, with α and γ phosphates; the structure of ATPase GET3, (PDB 2WOJ, chain B [99]). **B**. Asn finger in myosin II from ***Dictyostelium discoideum*** (PDB 1W9J). The The Asn^S-4^ finger is provided by the Switch I motif which is shown in yellow. **C.** Na^+^ ion as a stimulator in dynamin, (PDB 2X2E [100]). The switch I/K-loop motif is shown in lilac. Notably, a Gly residue coordinating Na^+^ is also contacting the O^2A^ atom of GDP. **D.** The LSGGQ motif in a structure of ABC-ATPase MBP-Maltose transporter MalK, (PDB 3PUW, chain A [93]). While Ser and Gly residues are coordinating γ-phosphate via the backbone HN atom and the OG atom of the conserved serine residue, the Gln residue is contacting the O2’ atom of GDP.

We have found that more than half of analyzed Mg-NTP complexes (60%) have none of the inspected residue types within the 4Å radius around oxygen atoms. Those are structures of P-loop NTPases that were crystallized in the absence of their activating partners or are stimulated by moieties other than Asn/Arg/Lys (e.g. a monovalent cation or the signature motif of ABC-ATPases, see below).

In the remaining 1380 catalytic sites of P-loop NTPases, at least one Arg, Asn, or Lys residue (other than the reference Lys residue of the Walker A motif) was found in the proximity of the phosphate chain and categorized as a stimulator. The analysis of interactions between Arg, Lys, and Asn fingers and the phosphate chains has revealed several distinct types of configurations, which hereafter are called “stimulatory patterns”. The Table S2 shows how many proteins were assigned to each stimulatory pattern.

Our provisional screening of structural information on different classes of P-loop NTPases has identified Arg, Lys and Asn residues, monovalent cations, and stimulatory polypeptide loops of ABC-NTPases as “main” stimulators, see [92] and the accompanying article [18]. Below we describe their interaction patterns.

In addition, we quantitatively evaluated the interaction patterns of Arg, Lys and Asn residues as main stimulators, as well as interactions of amino acid residues that serve as “auxiliary residues/stimulators”, see the Methods section and Table S1. The interactions of monovalent cations were quantified in our previous paper [70]. For ABC-NTPases, there was not much to quantify because their only two available TS-like structures were obtained for the same protein [93]. We saw that histidine residues may serve as stimulators in several NTPase families (as discussed in the Supplementary File 1 of the accompanying article [18]); however, for these families there are no TS-like structures so that we cannot state this for certain. We could not identify a glutamine residue as a dedicated stimulator in any structure, although Gln residues serve as auxiliary, W_cat_-coordinating ”fingers” in many P-loop NTPases, see Fig. 1C, 3, 5 and Table S1. We discuss this conundrum in the Discussion section.

##### 2.2.5.1. Stimulatory patterns of Arg fingers

Arginine fingers are the most widespread stimulatory moieties among P-loop NTPases. In an arginine side chain, the positive charge is distributed over three nitrogen atoms of the guanidinium group. In principle, each of these atoms can interact with the phosphate chain. Consequently, we observed a variety of interactions for the Arg fingers.

In most cases, the type of stimulatory pattern was assigned automatically based on the H-bond compatibility of distances between the NTP molecule or its analog and nearby Arg residue(s). Here we relied on Jeffrey who categorized H-bonds with donor-acceptor distances of 2.2-2.5 Å as “strong, mostly covalent”, those with distances of 2.5-3.2 Å as “moderate, mostly electrostatic”, and H-bonds of 3.2-4.0 Å as “weak, electrostatic” [83].

Due to inconsistencies in the atom numbering and differences among NTP analogs, we measured the distances from the Arg side chain nitrogen atoms to the nearest oxygen atom of α-phosphate (hereafter Oα) and the nearest oxygen of γ-phosphate (hereafter Oγ) or the corresponding atom in NTP analogs. Hereafter, for simplicity, we will use “γ-phosphate” both for γ-phosphate proper and for its analogs.

Several sets of criteria were applied consecutively, with each following criterion applied only to the cases that did not match any of the previous criteria:

1. If both distances NH1-Oα and NH1-Oγ do not exceed 3.2Å, the interaction type “NH1” was assigned, meaning that NH1 atom forms H-bonds with both α- and γ-phosphates. Similarly, “NH2” interaction type was assigned if both distances NH2-Oα and NH2-Oγ were less than 3.2Å.
2. If both distances NH1-Oα and NH1-Oγ do not exceed 4Å, whereas both distances NH2-Oα and NH2-Oγ are longer than 4Å, the interaction type “NH1 weak” was assigned, meaning that NH1 atom forms weak interactions with both α- and γ-phosphates. Analogous criteria were used to assign the “NH2 weak” interaction type.
3. If at least one of the distances NH1-Oγ and NH2-Oγ do not exceed 3.2Å, whereas both distances NH1-Oα and NH2-Oα are longer than 4Å, the interaction type “only gamma” was assigned. Similarly, if at least one of the distances NH1-Oγ and NH2-Oγ do not exceed 4Å, whereas both distances NH1-Oα and NH2-Oα are longer than 4Å, the interaction type “Only gamma weak” was assigned.
4. If all distances between NH1/NH2 atoms and the nearest oxygen (or fluorine) atoms of α- and γ-phosphates exceed 4Å, the Arg residue was considered not to be a stimulatory finger (interaction type “none”).

The remaining cases, which did not match any of these criteria, were inspected manually (see below). After the interaction types were assigned to all structures under investigation, the interaction types were attributed to particular stimulatory patterns and their frequencies were assessed, see Table S3.

Arg residues in the proximity of the phosphate chain were identified as stimulatory moieties in 981 cases. Majority of Arg fingers link α- and γ-phosphates by their NH1 or NH2 groups and fall into NH1, ”NH1 weak”, NH2, “NH2 weak” interaction types, which together are grouped into the stimulatory pattern “AG” seen in the case of 63% of all identified Arg fingers (Tables S1, S2, Fig. 6A-C). Among the structures with TS analogs, the fraction of this stimulatory pattern reaches 94%. In contrast, in complexes with ATP or GTP molecules, only 56% of interactions could be categorized in this way.

In most remaining structures, Arg fingers show interaction types “only gamma”/”only gamma weak” and interact only with oxygen atom(s) of γ-phosphate or their analogs (Fig. 6D, stimulatory pattern “G”). This stimulatory pattern was identified in 39% of complexes with ATP/GTP, 47% of complexes with nonhydrolyzable NTP analogs and only 6% of complexes with TS analogs (Table S2).

The remaining 33 complexes, which account for 3% of all Arg fingers, did not match any of these patterns. In these 33 cases, one NH1/NH2 atom of the Arg residue forms an H-bond with α-phosphate, whereas the other NH2/NH1 atom forms another H-bond with γ-phosphate. Further, we refer to such Y-shaped interactions as “Y-interactions” or “Y-patterns”. Since such Y-interactions are seen only in a small fraction of catalytic sites, we had inspected each of these sites manually; the results of this inspection are presented in the Supplementary File 1. As argued and illustrated in Supplementary File 1, there are reasons to consider all cases of Y-interactions as structure determination/crystallization artifacts of diverse nature.

Since the guanidinium group of Arg residues can donate several H-bonds, further H-bonds are seen between amino groups of the Arg finger and the oxygen atoms of the γ-phosphate (or its mimicking group). There are two types of such additional bonds: formed by the NE atom and formed by NH1/NH2 groups not involved in the main stimulatory interaction, as exemplified by Fig. 6.

The NE atoms of Arg fingers are often located at the H-bond distances from the γ-phosphate. Such interactions are documented for 10% of all Arg fingers, both for those Arg fingers that interact only with the γ-phosphate (Fig. 6D), and for those fingers that coordinate both α- and γ-phosphate with the NH2 atom (Fig. 6A).

An additional H-bond can be formed also by an NH1/NH2 atom which is not involved in the main stimulatory interaction. Usually, this occurs when one NH1/NH2 atom coordinates both α- and γ-phosphates; in 51% of such complexes, the other atom (NH2/NH1 correspondingly) forms a H-bond with γ-phosphate (Fig. 6B, C). This interaction is particularly common in complexes with TS analogs (77%). Finally, when the Arg finger interacts only with γ-phosphate, it can accept H-bonds both from NH1 and NH2 atoms, as observed in 13% of complexes with such interaction pattern (Table S2). In these cases, the longer H-bond was categorized as the “auxiliary” interaction, see Tables S1 and S2 for the complete data set.

Overall, the NH2/NH1 groups of Arg fingers that interact only with γ-phosphate or its analog are often assisted – in 40% of such complexes additional bonds are provided by the second NH1/NH2 atom, NE atom, or an additional Arg/Lys finger (Fig. 6D). In this case, one can speak about the stimulatory pattern G_multi_. The Arg fingers in the AG position can also receive assistance (Fig 6C). For example, in the FtsK DNA translocase structure (PDB ID 6T8B, chain C [97]), the Arg residue interacts with α- and γ-phosphates via the NH2 atom, while the Lys finger reaches the γ-phosphate. Arg fingers reaching only γ-phosphate often contact residues involved in the coordination of W_cat_ (see Fig. 6C, D).

##### 2.2.5.2. Stimulatory patterns of Lys fingers

Lys residues were assumed to be present in AG site if both NZ-Oα and NZ-Oγ distances are shorter than 4Å (stimulatory pattern “AG” see Fig. 7A and Table S1) and to interact only with γ-phosphate if only the second distance met the criteria (stimulatory pattern “G” in Table S1). Otherwise, no interaction with a Lys finger was presumed (pattern “None”), see Table S1.

Lys fingers were identified in 141 structures. One typical pattern is with the NZ atom of Lys interacting with both α- and γ-phosphates, similarly to a K^+^ ion in K^+^-dependent P-loop NTPases, cf Fig. 1D and [70]. Although a Lys finger interacts both with α- and γ-phosphates in 22% of all cases (we categorize these cases as stimulatory patterns AG, Fig. 7A), the fraction of such interactions was as high as 84% in complexes with TS analogs (Tables S1, S2). When the NZ atom of Lys interacts only with the γ-phosphate (pattern “G”), another Arg residue is also often involved in the interaction with the γ-phosphate (in 78% of cases, see Fig. 6D). Six of inspected structures had Lys finger coordinating both α- and γ-phosphate and an additional Arg residue in the proximity of γ-phosphate. All these structures are subunits of the Large T antigen (PDB ID 1SVM, in complex with ATP [98]), see Table S1 for details.

##### 2.2.5.3. Interaction patterns of Asn fingers

Asn residues were classified as stimulatory fingers when both ND2-Oα and ND2-Oγ distances were shorter than 4Å (stimulatory pattern “AG”, see Fig. 7B, in Table S1). The Asn residues were found to be in contact with both α- and γ-phosphate groups in 67 complexes. All these structures belong to myosin or kinesin families (PF00063, PF00225).

More common are auxiliary Asn residues, which were found in the proximity of γ-phosphates in 248 catalytic sites, in addition to the “main” stimulators, these auxiliary Asn residues are indicated in Table S1.

##### 2.2.5.4. Summary on quantitative analysis of stimulatory interactions of Arg, Lys, and Asn fingers in P-loop NTPases

As summarized in Tables S1, S2 and shown in Fig. 6 and 7A, B, most of the analyzed P-loop NTPase complexes with stimulatory Arg, Lys, or Asn residue(s) positioned next to the triphosphate chain (or its structural analog) possess either a residue providing an amino group to interact with both α- and γ-phosphates (stimulatory pattern AG, 56,6% with Y-interactions), or (2) an amino-group-providing residue(s) forming multiple bonds with γ-phosphate (stimulatory pattern G_multi_, 25,6%). In the rest of the cases (17,8%), only one amino group is contacting γ-phosphate (stimulatory pattern G_lone_).

In the case of TS analogs, the fraction of catalytic sites with stimulatory pattern AG is remarkably high: all structures with ADP:VO_4_^3-^ and NDP:MgF_3_^-^ that possess any kind of stimulatory residue have it interacting with both α- and γ-phosphates. The same interaction is observed in 75% of NDP:AlF_4_^-^ complexes whereas 22% of them display the stimulatory pattern G_multi_. Only 3% of TS analogs show stimulatory pattern G_lone_ with a single amino group contacting γ-phosphate.

The complete data on diverse kinds of auxiliary residues additionally stabilizing the negatively charged γ-phosphate and/or W_cat_ in all the studied structures are presented in Tables S1 and S2.

##### 2.2.5.5. Stimulation by monovalent cations

In at least two clades of P-loop NTPases, monovalent cations serve as stimulators, as already elucidated in a separate article [70]. First, many TRAFAC class GTPases are stimulated by K^+^- ions; the formation of a cation-binding site requires a particular positioning of the Switch I loop (dubbed K-loop in K^+^-dependent NTPases [27]), which is achieved either by the specific interaction of the P-loop domain with its activating partner(s) - protein and/or RNA molecules [70], or can be induced by binding of a TS analog, such as GDP:AlF_4_^-^ in Fig. 1D. The K^+^ ion in MnmE GTPase is coordinated by O^2A^, O^3B^, and O^3G^ atoms of the triphosphate chain, two **CO** groups of the K-loop, and the side chain of Asn^K-3^ (Fig. 1F, S2). [27, 33]). In the unique eukaryotic protein family of dynamins, the NTP hydrolysis can be stimulated by either K^+^ or Na^+^ ions, see Fig. 7C [100]. Here a Na^+^ or a K^+^ ion interacts only with the O^3B^ and O^3G^ atoms but does not reach the O^2A^ atom, see [70] for further details.

Second, in archaeal and eukaryotic RadA/Rad51-like recombinases of the RecA/F_1_ class, the positions that are taken by terminal groups of stimulatory Lys/Arg residues in other proteins of this class are occupied by two K^+^ ions, which might interact either only with γ-phosphate, or with the γ-phosphate and W_cat_, as could be inferred from available structures with substrate analogs bound, see Fig. 6D in the accompanying article [18] and [70, 101, 102]. No TS-like structures are available for these K^+^-dependent enzymes.

##### 2.2.5.6. Stimulatory interactions in ABC-NTPases

ABC (*ATP-binding cassette*) NTPases are multidomain proteins that usually operate as homo- or heterodimers [103]. Members of the ABC class make several families named alphabetically from A to I [104]. Most of these families contain members that possess transmembrane domains and operate as genuine ATP-driven membrane transporters where the P-loop domains hydrolyze ATP. However, the members of ABCE and ABCF families have no transmembrane domain(s) [105].

As discussed in more detail in the accompanying article [18], activation of ABC-transporter ATPases (ABC-ATPases) is triggered by the transported substrate and accompanied by dimerization of P-loop domains. Upon dimerization, each monomer, instead of an amino acid “finger”, inserts a whole signature motif LSGGQ into the catalytic pocket of the other monomer (Fig. 7D). Some soluble ABC-NTPases have a non-canonical signature motif (e.g. CSAGQ in Rad50 [106] and xSTFx in MutS [107]). In some cases, the last residue of the motif is Glu or even Trp [108]. Thus, the serine residue is the most conserved member of the signature motif.

Two structures with ADP:VO_4_ and ADP:AlF_4_^-^ as TS analogs bound are available for the maltose transporter complex [93], see Fig. 7D here and Fig. 5D in the accompanying article [18]. In both structures, the side chain of serine and the backbone **HN** of the second glycine residue of the signature motif L<u>S</u>G<u>G</u>Q interact with the O^3G^ atom of γ-phosphate. The side chain of serine is located between the α- and γ-phosphates, approximately in the position of the Na^+^ ion in dynamin-like proteins, cf Fig. 7C with Fig. 7D.

In sum, P-loop NTPases exhibit many various stimulatory patterns (see Fig.1-7 and Tables S1, S2) involving diverse atoms of stimulators, different phosphate groups and, in some cases, also the W_cat_ molecules (Fig. 1C). These interactions can play an instrumental role both in twisting the γ-phosphate group upon catalytic transition and, as discussed in more details in the accompanying paper [18], in constricting the catalytic site and the catalytic proton transfer.

## 3. Discussion

Here we report the results of a comparative structural analysis of 3136 catalytic sites of P-loop NTPases with nucleoside triphosphates or their analogs bound. The aim of the analysis was to find common features in their stimulatory patterns and to use this information for elucidating the general mechanism of these enzymes.

Our comparative structural analysis with emphasis on TS-like structures showed that highly diverse stimulatory moieties interact with the triphosphate chain only in two distinct ways that may complement each other, as discussed below.

### 3.1. Linking of α-and γ-phosphates by the stimulator

In most classes of P-loop NTPases, at least one stimulator, provided either by the same P-loop domain or by another domain/protein, gets inserted between α-and γ-phosphates, which implies the possibility of simultaneous interaction with O^2A^ and O^3G^ atoms (stimulatory pattern AG). As seen in Fig. 1, 3-7, in many such cases, the stimulator in the AG site is complemented by a second auxiliary stimulator (finger) that interacts with γ-phosphate, see also Tables S1, S2. The AG pattern was observed in 56.6% of cases.

Simultaneous interaction of a single amino group or a K^+^ ion with O^2A^ and O^3G^ atoms is possible either if the triphosphate chain bends (as it is observed with ATP or GTP molecules in water in the presence of Na^+^ ions [70]), or when the γ-phosphate group twists. The bending of the triphosphate chain of a P-loop-bound NTP molecule is not possible because of its multiple bonds with the protein, see Fig. 1C-D, the accompanying article [18], and [70]. At the same time, the examination of available structures revealed several cases which can be considered as evidence of γ-phosphate twist in the TS (see Fig. 4, 5), in support of our earlier predictions from MD simulations [70].

Quantification of the interaction types for computationally analyzed catalytic sites (Tables S1, S2) showed that the AG stimulatory pattern dramatically prevailed in the structures containing TS analogs. This pattern was observed in all structures with ADP:VO_4_^3-^ and NDP:MgF_3_^-^ and in 75% of NDP:AlF_4_^-^-containing structures. The fractions of this stimulatory pattern in structures containing ATP, GTP, or their non-hydrolyzable analogs are smaller (Table S2). In many cases, the Arg residue is in the AG-site in the presence of a TS analog but “outside” when the substrate or its analog are bound. Apparently, catalytic sites constrict additionally in the transition state; we elaborate on this point in the accompanying article [18].

The notable feature is the apparent scarcity – if not complete absence – of Y-patterns with NH1 and NH2 groups of an Arg finger separately interacting with α- and γ-phosphates, respectively (see Table S1). The Y-pattern is not observed in a single structure with a bound TS analog, and it is such structures that enable us to judge with certainty the stimulatory pattern in a particular ATPase. Our analysis has shown that the few structures with the Y-pattern are likely to be artifacts either of crystallization or of structure determination, as substantiated in the Supplementary File 1.

Outside of P-loop NTPases, however, the Y-pattern of Arg interaction is very common, especially in protein-DNA complexes, where one Arg residue often donates its NH1 and NH2 groups to neighboring phosphate groups of the DNA backbone [109, 110]. In the case of P-loop NTPases, however, a Y-linked Arg residue would fix the O^2A^ and O^3G^ atoms approx. 6.1 Å apart and prevent the twist of γ-phosphate. The apparent absence of Y-patterns in the examined structures of P-loop NTPases can be seen as further evidence in favor of the γ-phosphate twist as the key precatalytic configuration change in P-loop NTPases.

Earlier, our MD simulations showed that γ-phosphate, after its twisting by the stimulatory K^+^ ion, was stabilized by a new H-bond between HN^K-3^ and O^2G^ [70]. Our structural analysis strongly indicates that the counter-clockwise twist of γ-phosphate correlates with the formation of a new H-bond between the backbone of the P-loop and O^2G^ indeed. Specifically, H-bond-compatible distances between **HN**^K-3^ and the nearest oxygen/fluorine atom are seen in most structures with TS analogs bound (Fig. 2, Table S1). Twisted γ-phosphates and H-bond compatible **HN**^K-3^ - O^2G^ distances are also seen in diverse NTP-containing proteins which were crystalized after being trapped in their pre-transition configurations (because of mutated W_cat_- stabilizing residues and/or their impaired interaction with W_cat_, see Fig. 3A-C, Table S1).

Hence, linking the O^2A^ and O^3G^ atoms by the stimulator, independently on whether it is an Arg, Lys, Asn, residue or a K^+^ ion, causes a counter-clockwise twist of γ-phosphate that correlates with the formation of a new H-bond between the O^2G^ atom (or its counterpart in NTP analogs) and the backbone HN^K-3^ group of the P-loop.

### 3.2. Interaction of the stimulator only with γ-phosphate

In the remaining 43,3% of P-loop NTPase structures with determined stimulatory pattern, the stimulators interact only with γ-phosphate (stimulatory pattern G). In most such structures (25,6%), γ-phosphate is involved in several interactions with distinct amino groups of an Arg finger and/or with auxiliary stimulators (stimulatory pattern G_multi_), see Fig. 6D, 7D and [111].

In 17.7% of structures, our computational analysis has reported only one H-bond between the stimulatory residue and γ-phosphate (stimulatory pattern G_lone_). However, in many cases, the crystal structure does not contain all the partners involved in the activation, or additional stimulatory fingers are present but are too remote because of crystallization artifacts. In fact, many catalytic sites exhibiting a G_lone_ pattern have counterparts with a richer network of H-bonds around γ-phosphate in other subunits of homooligomers of the same protein (47.6%) or in proteins belonging to the same Pfam domain (75%). Consequently, the value of 17.7% should be at least halved.

While linking of O^2A^ and O^3G^ atoms by a stimulator enforces a counterclockwise twist of γ-phosphate, it is not clear yet, what conformational changes are caused by stimulators that interact only with γ-phosphate. There is indirect evidence that γ-phosphate may be twisted clockwise in RecA NTPases, see [112, 113]. Also, in the structure of the ABC ATPase of the maltose transporter (PDB ID 3PUW, [93]) the AlF_4_^-^ moiety is slightly twisted clockwise (Fig. 7D).

Even interacting only with γ-phosphate, the stimulator is often located between α- and γ-phosphates, as in dynamins (Fig. 7C) or ABC transporters (Fig. 7D) and connected to the “head” of the NTP molecule. For instance, the Na^+^ ion in dynamins, while not reaching the α-phosphate directly, is connected to it via two noncovalent bonds (Fig. 7C). The signature motif of ABC-NTPases is H-bonded via conserved Ser and Gln residues to the O2’ atom of the ribose (Fig. 7D). Such connectivity may strengthen the mechanistic impact on γ-phosphate.

In sum, our structural analysis shows that in all cases where stimulators reach only γ-phosphate, these stimulators, independently on whether they are Arg or Lys residues, K^+^ or Na^+^ ions, or signature motifs of ABC-ATPases, are in position to twist or pull the γ-phosphate group.

### 3.3. Role of mechanistic bonding in the common stimulation mechanism of P-loop NTPases

Looking together at all types of identified stimulatory patterns provides some additional clues about the mechanisms of hydrolysis stimulation. Without challenging the previously proposed tentative stimulatory effects as referred to in the Introduction section, our structure analysis indicates that none of so far suggested mechanisms is common to all P-loop NTPases. Indeed, it is beyond doubt that the positive charges of Arg/Lys fingers or K^+^/Na^+^ ions, by interacting with oxygen atoms of γ-phosphate, would make the P^G^ atom more prone to the nucleophilic attack, as suggested by Warshel and colleagues [60–62] and as calculated by Rudack et al. [67]. The positive charge of stimulators could also compensate for the negative charge that develops at β-phosphate upon the breakaway of γ-phosphate [49, 66]. Nevertheless, the absence of a positive charge on the stimulatory signature motifs of ABC ATPases (Fig. 7D) does not stop them from triggering ATP hydrolysis. Also, expelling water molecules out of the catalytic pocket by stimulatory Arg fingers may provide an entropic gain, as suggested by Kotting and collegues [63]. However, such effects are hardly to be expected when K^+^ or Na^+^ ions act as stimulators and immobilize water molecules in the catalytic pocket (Fig. 1D, 7C). Reorientation of the W_cat_ molecule into the attack position and its polarization, as suggested by Jin and colleagues [44, 64, 65], can be realized by those Arg or Lys fingers that reach W_cat_ (Fig. 6D), but not by most other stimulators.

Our analysis points to the importance of mechanistic interaction of stimulators with the γ-phosphate group, which is the only feature shared by all inspected stimulators. Importance of this interaction is exemplified by NTPases that are stimulated by moieties with only a minute positive charge. These are ABC-NTPases, where γ-phosphate binds the signature LSGG[Q/E] motif via the side chain of its serine residue and the backbone **HN** of the second glycine residue, see Fig. 7D and [93]. Other examples are the kinesin and myosin families, where the Asn finger inserts its NH group between α- and γ-phosphates, see Fig. 7B and [114–116]. It is unlikely that small partial electric charges of Ser or Asn side chains could be decisive for catalysis in these cases; rather, their mechanistic H-bonding to the O^3G^ atom appears to be the key.

The mechanistic nature of the stimulatory interaction is consistent with the predominance of Arg residues as stimulators (Table S1-S3). First, a guanidinium group could donate up to three H-bonds for interaction with the oxygens of triphosphate. Second, the strength of H-bonds between guanidinium groups and phosphate anions has been shown to be comparable to that of covalent bonds [117–119].

The importance of mechanistic binding rationalizes the preference for multiple stimulatory fingers (Table S2), as well as the choice of the stimulatory signature motif by omnipresent ABC ATPases. This motif is electrically neutral but donates several H-bonds to the O^3G^ atom.

Consequently, the common denominator of stimulatory patterns in diverse P-loop NTPases is the mechanistic interaction of stimulators with the γ-phosphate group; this interaction is observed in all analyzed TS-like structures (Fig. 1C,D, 3-7, Table S1, S2) and can be inferred from many other structures, specifically those that can be related to post-transition states, see [113] and the accompanying paper [18].

Mechanistic interaction with the γ-phosphate group may promote hydrolysis in different ways; for instance, it may destabilize the O^3B^ – P^G^ bond and/or make the triphosphate chain almost fully eclipsed, and/or facilitate the inversion of γ-phosphate, see the discussion in [70] and the accompanying article [18]. Notably, any rotation of γ-phosphate inevitably disturbs the coordination sphere of Mg^2+^, since the O^1G^ atom of γ-phosphate is one of the Mg^2+^ ligands (Fig. 1B-D). In P-loop NTPases, the O^1G^ atom is negatively charged, so that its displacement, by affecting the proton affinity of the other five Mg^2+^ ligands, may trigger the deprotonation of W_cat_. In more detail, we address this point in the accompanying article [18] where we further use comparative structure analysis to reconstruct those steps of the catalytic transition that follow the interaction with the stimulators.

### 3.4. The puzzling absence of glutamine residues as stimulators

Glutamine residues are involved in stabilization of W_cat_ in many P-loop NTPases, see e.g. Fig. 1C, 3, 4, 6A,B and the accompanying paper [18]. Still, Gln residues, unlike Asn residues (Fig. 7B), could not be identified as actual stimulators in any of catalytic site structures. Notably, however, Gln residues occupy stimulator-like positions in non-catalytic sites of F_1_-ATPases (see a plethora of entries in Table S1). These sites, however, are non-functional (see also Supplementary File 1). It is tempting to speculate that the excessive, as compared to Asn, flexibility of the glutamate side chain prevents it from twisting γ-phosphate.

### 3.5. Geometry of the AlF_3_ moiety in the NDP:AlF_3_ complexes

As a side result, we determined the geometry of an Mn^2+^:ADP:AlF_3_ complex in a P-loop ATPase (Fig. 4) and thus offered a solution to the long-standing controversy as to whether NDP:AlF_3_ complexes could form in the catalytic site of P-loop NTPases under appropriate conditions or whether all such complexes are misassigned NDP:MgF_3_^-^ complexes, see [35, 42, 44, 58]. The Mn^2+^:ADP:AlF_3_ complex was crystallized in the absence of Mg and therefore could not contain a MgF_3_^-^ moiety. The identified AlF_3_ moiety has substrate-like geometry of the AlF_3_ group, unlike the planar, TS-like MgF_3_^-^ moieties. This non-planarity may help to discriminate such complexes from NDP:MgF_3_^-^ complexes in earlier obtained crystal structures. Hence, it is realistic to sort out the P-loop-bound NDP:AlF_3_ complexes, as assigned in the PDB, into misassigned NDP:MgF_3_^-^ complexes and genuine NDP:AlF_3_ complexes.

### 3.6. Unwelcome effects of AlF_4_^—^ binding

Quite unexpectedly, we found that the AlF_4_^—^ moieties are bound in some catalytic sites in such a way that two fluorine atoms interact with the Mg^2+^ ion. In most of these cases, the coordination bond between Mg^2+^ ion and its ligand #4, [Ser/Thr]^K+1^, is lost, leading to a distortion of the catalytic site (Fig. 5, Tables S1, S3). This finding is alarming because fluoride complexes are deservedly vaunted as powerful TS analogues, and structures containing them are commonly interpreted as TS-like [35, 40, 44]. Still, our data show that the integrity of Mg^2+^ coordination in the presence of NDP: AlF_4_^—^ should be evaluated separately for each enzyme structure before linking the corresponding structural data to the catalytic mechanism. Notably, NDP:AlF_4_^—^ complexes are also used as TS analogues in other enzyme superfamilies [35, 44]. It is important to check whether a similar distortion of catalytic sites by NDP:AlF_4_^—^ complexes is possible in enzymes other than P-loop NTPases.

Although this discovery was rather unpleasant as such, it provided some useful information. The “wrong” binding of AlF_4_^—^ was accompanied by its counterclockwise rotation as compared to “properly bound” AlF_4_^—^ moieties, whereby the interactions of the fluoride atoms with Lys^WA^ and the stimulator were preserved. It could be inferred that the ligands of γ-phosphate oxygens are adapted to the twist of the γ-phosphate group.

### 3.7. Conclusions and a Flashback

Here, we performed the first, to the best of our knowledge, comparative structural analysis of the interactions between the stimulatory moieties and substrate molecules in P-loop NTPases of the major classes. After screening over 3,100 available structures of catalytic sites with bound Mg-NTPs or their analogs, we found that seemingly different interactions between completely distinct stimulatory moieties (Arg/Asn/Lys residues, or K^+^/Na^+^ ions, or LSGGQ/E motifs) come down to only two stimulatory patterns. In most cases, at least one stimulator links the O^2A^ atom of α-phosphate with the O^3G^ atom of γ-phosphate, which requires counterclockwise twist of the γ-phosphate group. In other cases, stimulators only interact with the γ-phosphate group. In general, the only shared feature of all the identified stimulators seems to be the ability to enter a mechanistic interaction with the γ-phosphate group, which may enable its twist/rotation.

We provide the relevant outlook on the future studies of P-loop NTPases at the end of the accompanying paper [18]. Here, instead, we would like to give a flashback.

In 1998, Wittinghofer and colleagues published a seminal review with the telling title “GTPase-activating proteins: helping hands to complement an active site,” where they viewed a variety of activating molecules as hands with “fingers” [31]. In this context, our data show that removing γ-phosphate from an NTP molecule resembles plucking an apple from a tree - the “fingers” seem to need to twist the γ-phosphate group before they can “rip it off”.

## 5. Methods

Structures were selected among those PDB entries that matched the following criteria: (1) the entry is assigned to InterPro record IPR027417; (2) contains an ATP/GTP molecule, or non-hydrolyzable analog of NTP, or a transition-state analog; (3) contains at least one Mg^2+^, Mn^2+^ or Ca^2+^ ion. We considered X-ray structures as well as cryo-EM and NMR structures. This search yielded 1474 structures with 3666 catalytic sites in them. For the NMR structures, only the first frame containing an NTP analog bound to the P-loop region was included. For X-ray and cryo-EM structures an additional criterion was applied: structures with resolution less than 5 Å were excluded. In the case of structures containing multiple subunits or multiple copies of the same protein, each interaction between a protein and an NTP-like molecule was treated separately, as an individual complex. The pipeline that was applied to each complex is depicted in Fig. 8.

**Figure 8.**
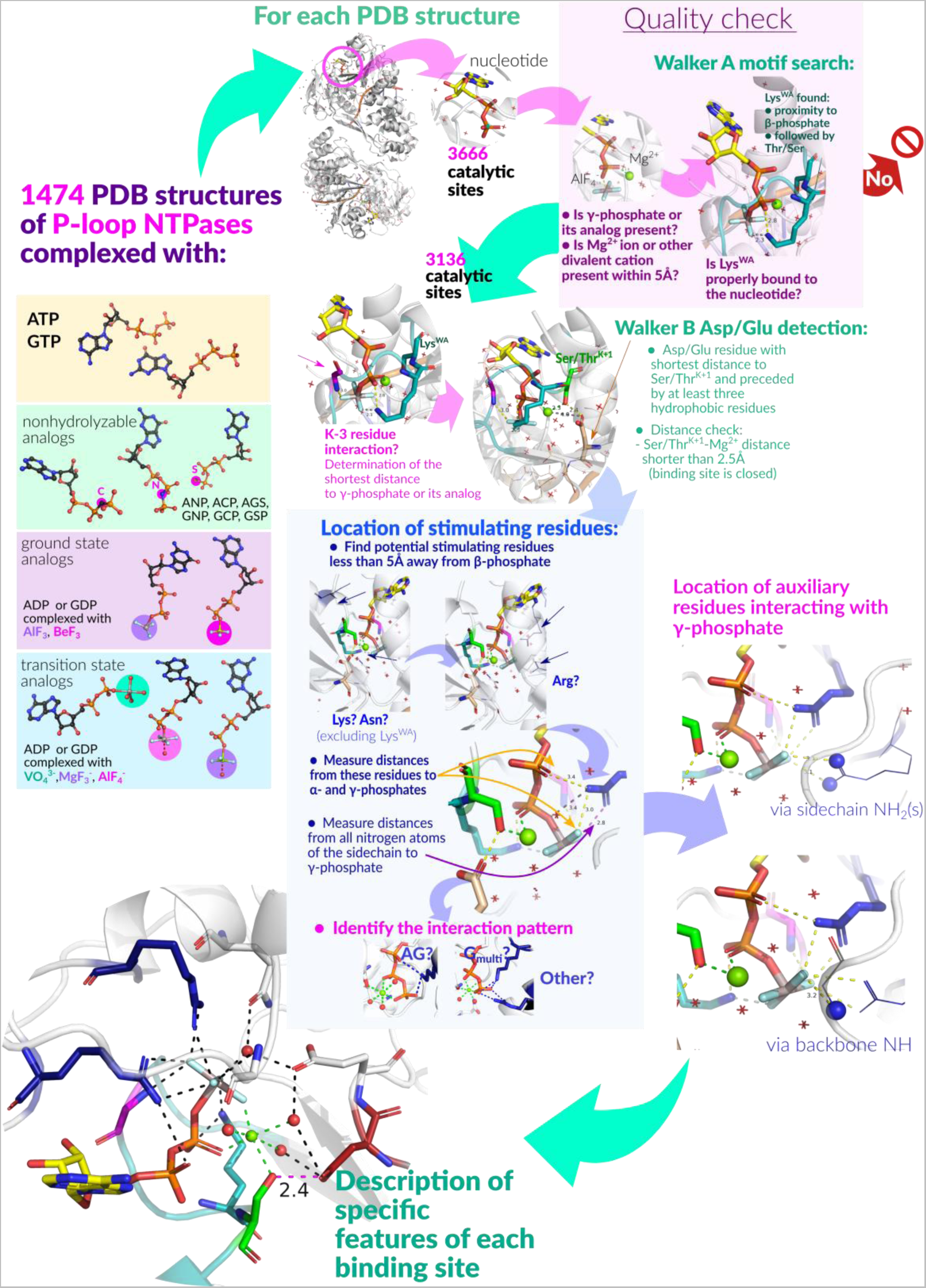
Automated comparative structural analysis of P-loop NTPases. The examination pipeline, as applied to the structure of each selected nucleotide-binding site, is shown. We checked for (i) the presence of the HN^K-3^ – O^2G^ bond, (ii) the length of the [Asp/Glu]^WB^-[Ser/Thr]^K+1^ H-bond and (iii) the type of interaction of stimulatory residues. Scripts used to download and analyze the structures are available at github.com/servalli/pyploop

To identify structures with a substrate or its analog bound to the P-loop motif, we applied the following filter: an NTP(-like) molecule was considered bound to the P-loop if both Mg^2+^/Ca^2+^/Mn^2+^ cation and the NZ atom of Lys^WA^ can be found within 5Å of the oxygen atoms of the β-phosphate. Furthermore, Lys^WA^ must be followed in the sequence by a Ser or Thr residue. Complexes with molecules annotated as nonhydrolyzable NTP analogs but lacking β- and/or γ-phosphate mimicking groups were not included in the analysis.

In total, we had analyzed 3136 complexes in 1383 structures with various substrates: ATP and GTP, nonhydrolyzable analogs of ATP and GTP, and ADP or GDP molecules associated with γ-phosphate-mimicking moieties (see Fig. 14).

In each catalytic site under consideration, the K-3 residue was identified by its position relative to the P-loop Lys residue. Distances (in Å) were measured from the **HN** of the K-3 residue to the nearest oxygen or fluorine atom of the *γ-*phosphate or its mimic. We accounted for differences in phosphate oxygen atoms numeration among studied structures.

Putative [Asp/Glu]^WB^ residue was identified as follows: distances from all Asp and Glu residues to [Ser/Thr]^K+1^ were measured, and the closest residue that was preceded by at least three non-ionizable residues (Glu, Asp, Ser, Thr, Tyr, Lys, Arg and His were considered as ionizable) was chosen as the partner of [Ser/Thr]^K+1^. Residues located further than 5Å were not considered.

We have also measured distances from [Ser/Thr]^K+1^ to Mg^2+^, to ensure the correct binding of the Mg^2+^ and general reliability of the structure resolution at the binding site (i.e, very long distance would indicate a disturbed catalytic site or resolution at the site that is insufficient for our purposes of comparative analysis), and from [Asp/Glu]^WB^ to Mg^2+^, to identify cases of direct coordination of Mg^2+^ by the acidic residue (short distances) or disassembled binding sites (long distances).

We had also inspected the presence of positively charged stimulatory residues near the phosphate chains in all complexes. To identify such residues, we considered Arg and Lys residues (excluding the P-loop Lys) nearest to the β-phosphate group oxygen atoms or its structural analogs. Distances were then measured from the guanidinium group nitrogen atoms (NE, NH1, NH2) of the Arg residue or from the NZ atom of the Lys residue to the closest oxygen atom of α-phosphate moiety and to the nearest fluorine or oxygen atom of γ-phosphate or its mimicking group. Similarly, we checked possible interactions of the phosphate chain with the ND2 atom of Asn. Possible additional interactions of γ-phosphate oxygens with N atoms of protein backbone and side chains in vicinity were also listed for each complex. For all these interactions the distance threshold was 4 Å.

In the systematic analysis of all available structures, the patterns of Arg finger binding were assigned automatically, based on the composition of H-bonds between the Arg residue and the substrate molecule. We had considered each pair of possible donor and acceptor atoms, where donors were NE/NH1/NH2 atoms of Arg residue (see Fig. 6) and the acceptors were oxygen/fluorine atoms of the substrate. The presence of an H-bond was inferred from the atomic distance: an H-bond was stated at distances less or equal 3.2Å, a weak H-bond stated at distances between 3.2 and 4Å, and no possibility of a H-bond was stated for distances over 4 Å [82, 120, 121]).

To assign the Arg finger types, several sets of criteria were applied consecutively, as described in the Results. Structures that did not fit any of the criteria were additionally inspected and Arg binding patterns assigned manually. After all the Arg fingers were categorized, the frequency of each interaction type was determined together for the automatically and manually assigned types.

The lysine residues were assumed to be present in AG site if both NZ-Oα and NZ-Oγ distances are shorter than 4Å (finger type “AG”), and to interact only with γ-phosphate if only the second distance met the criteria (finger type “G”). Otherwise, no interaction with a Lys finger was presumed (finger type “None”).

For each complex, AG site was interpreted as occupied by an Arg residue if the closest Arg residue was assigned interaction type “NH1”, “NH1 weak”, “NH2”, “NH2 weak” or “Y-TYPE”, or by a Lys residue if the closest Lys was assigned type “AG”.

All other Arg or Lys residues located within 4Å from γ-phosphate/γ-mimic were listed as present in “G-site”.

Proteins were assigned to major classes of P-loop NTPases according to membership in Pfam families that was determined from PDB-to-Pfam mapping (retrieved 10.10.2020 from ftp://ftp.ebi.ac.uk/pub/databases/Pfam/mappings/pdb_pfam_mapping.txt). Each chain was treated separately and only the first Pfam domain included in Pfam clan CL0023 (“P-loop_NTPase”) was used for the assignment. Since many Pfam domains describing P-loop NTPases were described before a coherent classification of P-loop NTPases was developed [7, 8], some domain names may not be accurate.

### Visualization

Structure superposition, manual distance measurements, manual inspection and structures visualization were performed by Mol* Viewer [122] and PyMol v 2.5.0 [123].

## Data availability

Descriptions of each binding site are available in Table S1. Scripts used to generate and annotate the data and quickly visualize selected sites listed in Table S1 are available from github.com/servalli/pyploop.

## 6. Acknowledgements

Very fruitful discussions with Drs. D.A. Cherepanov, M.Y. Galperin, A. V. Golovin, Y.Kalaidzidis, E.V. Koonin, B.H. Meier, A. Lupas, N. Voskoboynikova, T. Wiegand and M. Zereal are highly appreciated. The authors are thankful to Prof. H.-J. Steinhoff for useful suggestion on improving the manuscript. We also are thankful to Alexander Mulkidzhanyan for his help during the initial stage of the project. The research was supported by DFG, DAAD, and the Osnabrueck University (the EvoCell Program and Open Access Publishing Fund).

## Supplementary Materials

### Supplementary Figures

**Supplementary File 1.** Detailed consideration of structures with an atypical “Y-type”-like interaction pattern between the stimulatory Arg finger and the triphosphate chain (or its mimics in case of NTO analogs).

**Supplementary Table 1**. Results of the computational analysis of all available structures of the P-loop proteins in complex with Mg-NTPs or their analogs. The Excel table contains the list of all analyzed structures, together with residues, identified as the P-loop K and K-3 residues, Arg or Lys fingers; and distances from (1) the respective atoms of NTPs/their analogs to K-3 residues and Arg/Lys fingers, as well as (2) from [Asp/Glu]^WB^ to [Ser/Thr]^K+1^.

**Supplementary Table 2**. Relative occurrence of activation patterns for Arg, Lys, and Asn fingers.

**Supplementary Table 3**. Coordination of the Mg^2+^ ion in the AlF_4_^—^-containing structures of P-loop NTPases

## Supplementary Figures

**Figure S1.**
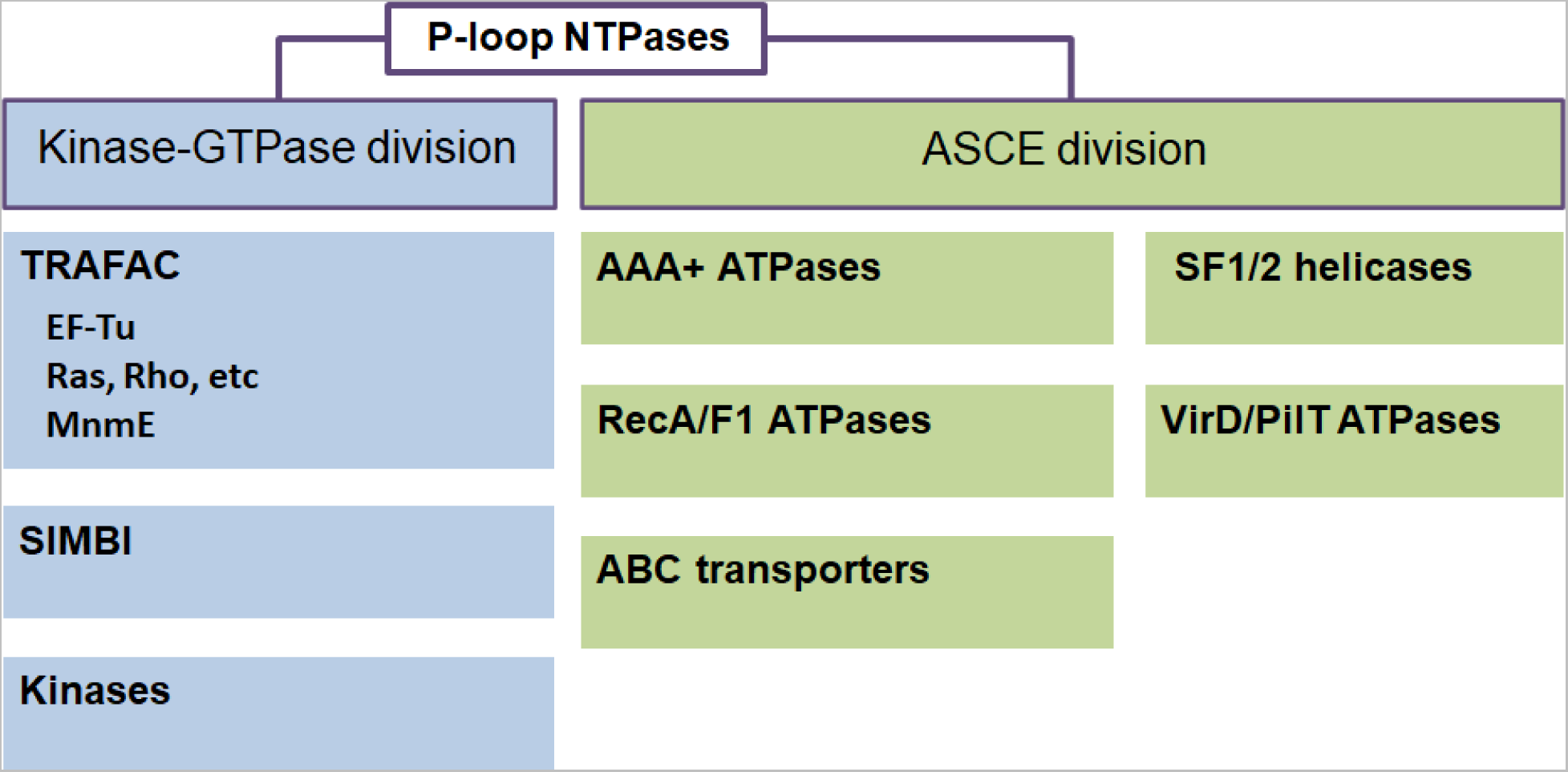
Major classes of P-loop fold NTPases according to [1–5].

**Figure S2.**
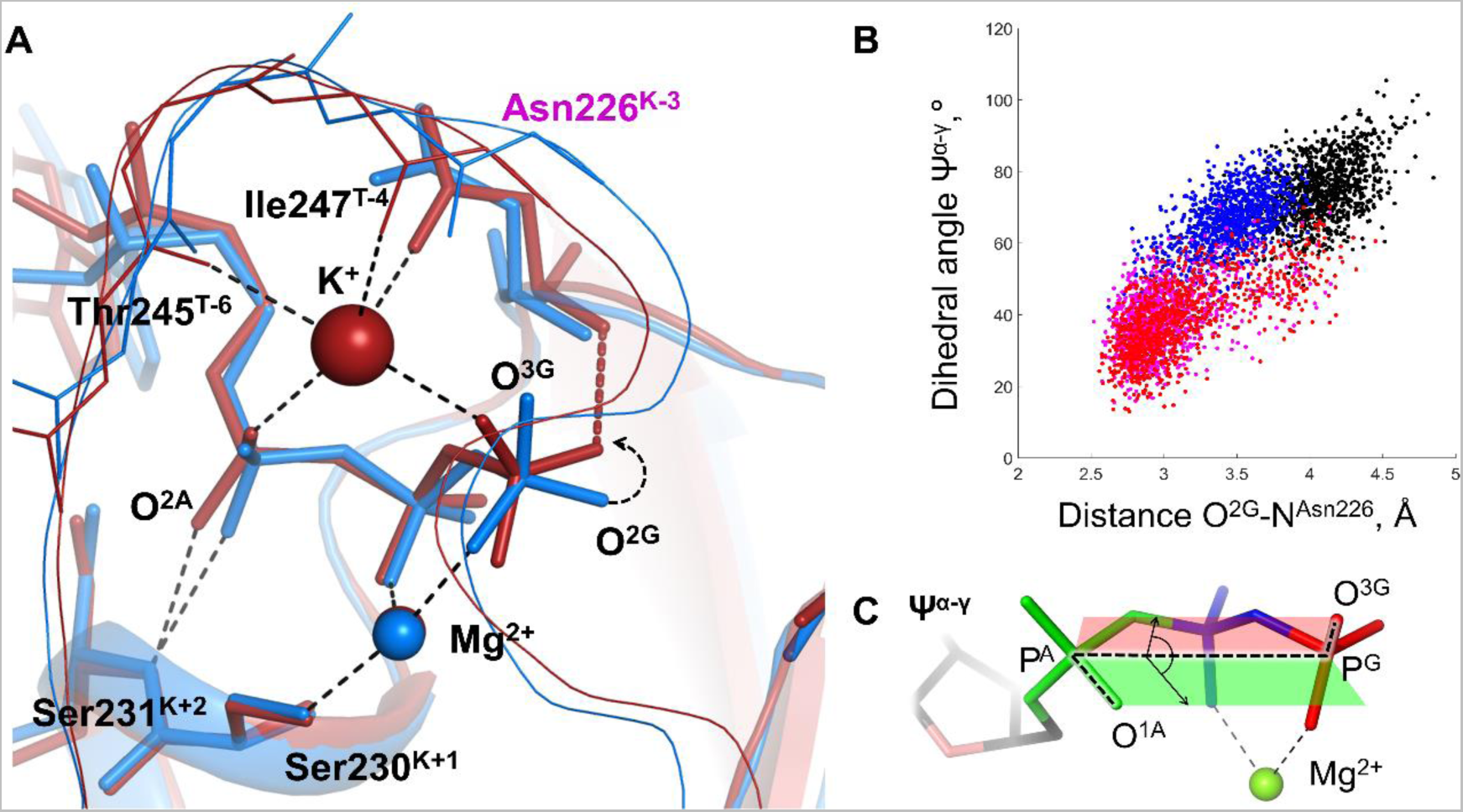
Molecular dynamics of the MnmE GTPase. (The figure is taken from [6] and modified). **A**. Superposition of the GTP-binding sites of the inactive, monomeric G-domain of MnmE (blue) and active, K^+^-bound G-domain in a dimer (red); the representative structures were sampled from 100 ns simulations as described in [6]. The protein backbones are shown as cartoons; GTP and surrounding amino acid residues are shown as sticks; Mg^2+^ and K^+^ ions are shown as spheres. Black dashed lines indicate hydrogen bonds and coordination bonds for cations that are present in both structures; the red dashed line indicates the H-bond between **NH^K-3^** and O^2G^ that is present only in the K^+^-containing dimer. **B**. Conformational space of GTP in different states of MnmE GTPase. Scatter plot of the Ψ^α-γ^ dihedral angle (Y-axis) against the distance between the O^2G^ atom and **NH** of Asn226^K-3^ (X-axis) as sampled from the MD simulations of three systems: (1) active dimer of G-domains with K^+^ ions bound (red and magenta for individual monomers); (2) monomeric G-domain of MnmE with the K^+^ ion replaced by a water molecule, blue; and (3) inactive monomer G-domain of MnmE without a full-fledged K-loop, black. **C**. The dihedral angle Ψ^α-γ^ in the phosphate chain of GTP as measured for the plot on panel B.

## Supplementary File 1

### Analysis of structures with Y-like interaction pattern of arginine fingers

Our analysis revealed only 33 complexes with Y-like interaction of an Arg finger with an NTP molecule or its analog (see also Table S1). The Y-pattern is not observed in a single structure with a bound TS analog, and it is such structures that enable us to judge with certainty the stimulatory pattern in a particular ATPase. Therefore, we inspected these 33 complexes manually.

**I).** Seventeen of these binding sites belong to the structures obtained via electron microscopy (EM), mostly with resolution worse than 3.5 Å. Eleven of these NTP binding site structures are of catalytically inactive ATP-binding sites of α-subunits of F_1_-type ATPases, from *Sus scrofa* (PDB 6J5J, chain A [1] and *Polytomella sp. Pringsheim 198.80* (all structures from [2]). In other 190 structures of non-catalytic sites, a Gln residue in K-3 position links the O^2A^ and O^3G^ atoms, see Table SF1 below. However, in these eleven structures, Gln^K-3^ does not reach the O^2A^ atom so that the “Y-type” Arg residue of the adjoining monomer takes the canonical “finger position” and enters a Y-interaction with O^2A^ and O^3G^. However, the high-resolution X-ray structures of the same non-catalytic sites reveal the amino group of the Gln^K-3^ residue in the AG position, as discussed in the main text. We do not know why these eleven structures show a different interaction than the rest 190 structures of noncatalytic sites. Anyhow, both the Gln “plug” in 190 structures, as well as the Y-interacting Arg in eleven deviating structures are fully compatible with the major task of non-catalytic ATP binding sites, which is <u>not</u> to catalyze ATP hydrolysis. The remaining six EM-derived complexes with Y-type interaction come from structures of oligomeric complexes where other subunits display a typical stimulatory interaction via a single NH_2_ group. Only one of these structures has a resolution better than 3.5 Å.
**II).** We manually inspected the remaining **15** binding sites with Y-pattern (as obtained by X-ray crystallography of **13** crystal structures) and found out that they can be attributed at least to one of the following five cases (see details for each binding site in Table SF1):

1) **Presence of other complexes of the same protein with other, more common stimulatory patterns, with only one NH_2_ group of an Arg finger interacting with both α- and γ-phosphates or stimulators interacting with only γ-phosphate.** Table SF1 contains five oligomeric structures where one protein subunit exhibits Y-like interaction of the Arg finger, whereas other subunits of the same protein have a different configuration of the catalytic site. No structures display two or more catalytic sites with a Y-like interaction for the same oligomeric protein. For some of the proteins with Y-pattern in Table SF1, there are many other structures either from the same (ten complexes) or different organisms (six complexes) that display a different binding pattern. For instance, the Y-pattern is seen in one catalytic site of the rotary ATP-synthase (PDB 3OEE, chain L). This binding pattern is not displayed either in the other catalytic sites of the same structure, or in 69 other catalytic sites from β -subunits with either ATP or its analogs bound and an Arg finger present (see Table SF1).
2) **Arg finger residue is listed as an outlier in wwPDB structure quality assessment reports.** In five cases, the Arg finger residues involved in Y-interactions were reported as outliers in regard to side chain geometry. Four of them are reported to possess a non-rotameric sidechain, while one residue is a bond angle outlier (NE-CZ-NH2 angle), see Table SF1. The reports also list too-close contacts; eight sites possess an Arg finger engaged in interatomic clashes either with other amino acid residues or with atoms of nucleotide moiety that are not expected to form H-bonds. In the structure of Guanine nucleotide-binding protein G (PDB 3FFA [3]) the sidechain atoms of Arg178 are even clashing with neighboring atoms of the same residue. Specifically, this set of cases contains the stimulatory Arg789 residue of H-RasGAP that is Y-linked with α- and γ-phosphates in the first prototypical structure of the H-Ras/RasGAP complex (PDB ID 1WQ1 [4]), which has been widely used for MD and QM/MM modeling. The PDB X-ray Structure Validation Report for this structure (accessible at https://www.rcsb.org/structure/1WQ1) indicates that the conformation of the side chain of the Arg789 finger contains at least one outlier for two of the geometric quality criteria; it is noted also that Arg789 has a non-rotameric side chain conformation, which may point to a crystallization artefact. Notably, in earlier reported structures of highly homologous Gα-proteins the Arg finger interacted with α- and γ-phosphates via a single NH_2_ group (PDB ID 1GFI, 1GIL [5]). Also in the all subsequently obtained structures of Ras-like GTPases crystallized with TS-analogs and cognate activators (with resolution < 2.0 Å, see, for instance, the high-resolution structures with PDB ID 1OW3 [6], 1TX4 [7], 3MSX, 5IRC [8]), only one NH2 group interacts both with α- and γ-phosphates. This prompts the suggestion that the Arg residue in the very first structure of the Ras/RasGAP complex (PDB ID 1WQ1 [4]) should have the same orientation. This suggestion is corroborated by MD simulations of the Ras/RasGAP which, after starting from the crystal structure, promptly yielded a conformation where a single NH_2_ group interacted with α- and γ-phosphates, see e.g. [9].
3) **The distances between the NH2 group and O^2A^, O^3G^ atoms that are too short for an H-bond (O..H-N distance less than 2.4 Å) are observed in five complexes, see Table SF1.**
4) **Electron densities (ED) are inconsistent with the positioning of the residue side chains.** We have manually evaluated electron density maps (2Fo-Fc) for all 15 sites, see examples in Fig SF1A-F. Four Arg fingers lack ED entirely (example: Fig SF1A, F), while in one site the sidechain is poorly fitted to the available density (Fig SF1E). In two cases electron density distribution is more consistent with the same NH_2_ group bonded both to α- and γ-phosphates (Fig SF1E, B). Six complexes display some electron density, but it is poorly resolved in the terminal region of Arg residue, thus not allowing determination of the exact location of the guanidinium group (examples: Fig SF1C, D).
5) **Optimized structures in PDB REDO depict a different configuration of the stimulator.** We also checked the structures with Y-pattern in the PDB REDO structure databank, which contains automatically optimizes crystallographic structure models [10]. In three cases, the Y-pattern is absent from the re-refined structure, in two other sites the Arg is still inserted in a Y-like manner, however, the distance between the respective NH_2_ group and the more distant phosphate group is shortened. It is worth noting that sidechains lacking electron density cannot be expected to shift considerably in a re-refined structure.

Finally, eight complexes with Y-like patterns belong to SF1/SF2 helicases (see Fig 1C in the main text for an example of a typical binding site in a SF1 helicase). Remarkably, most (105) other complexes of this class, including all the complexes with TS analogs bound, depict a single NH_2_ group interacting with α-and γ-phosphates, and in seven remaining cases Arg/Lys residue(s) contacts only γ-phosphate. Almost all these complexes harbor a second Arg residue contacting the γ-phosphate moiety. We would suggest that the catalytic site is not properly arranged in the absence of TS analog in these eight structures with Y pattern.

In sum, these findings indicate that a Y-like stimulatory pattern is unlikely to be inherent to P-loop fold NTPases; its presence in a few experimental structures might be due to poor resolution of particular residues or crystallization artefacts.

---

**Figure SF1.**
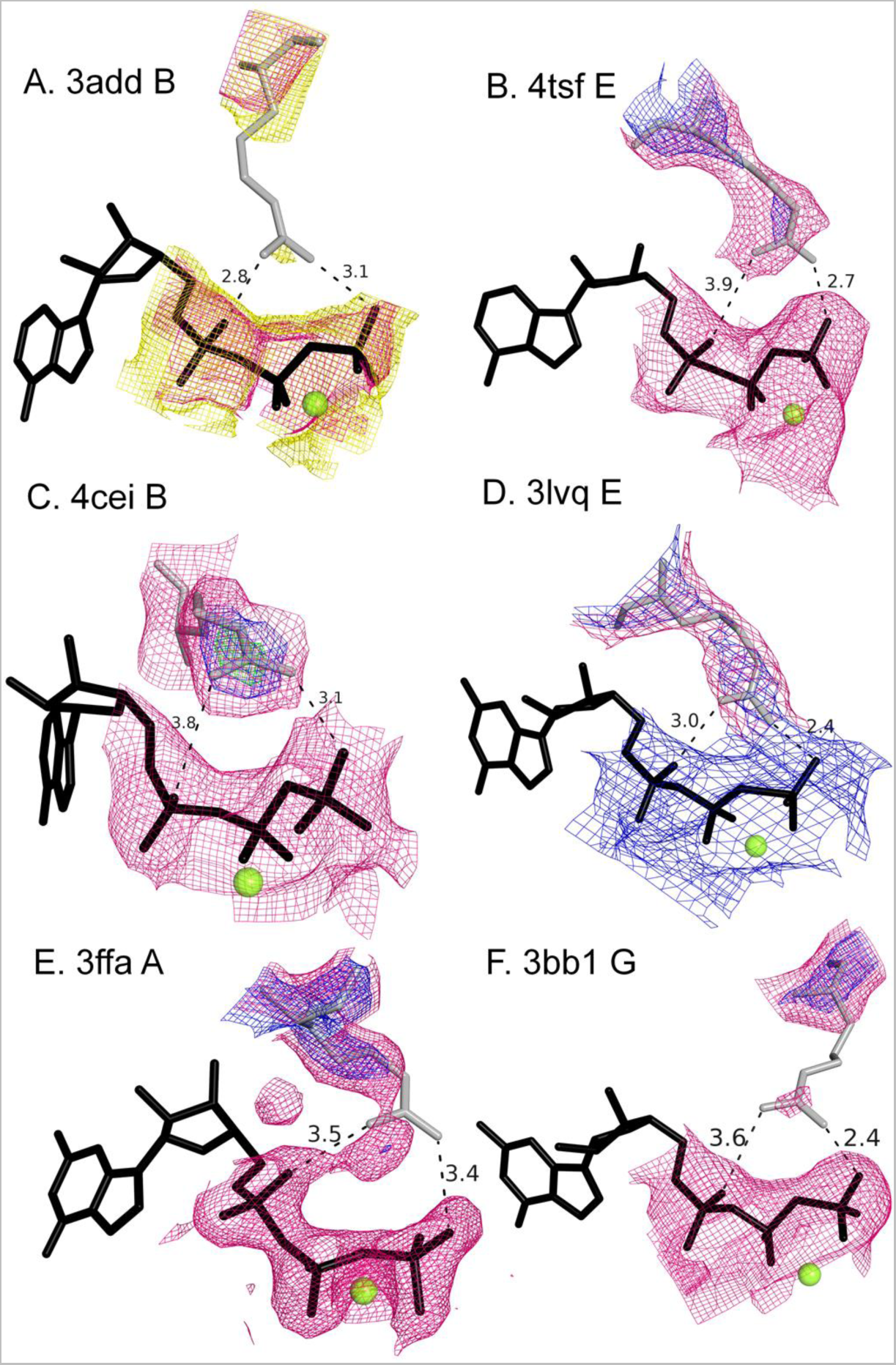
2Fo-Fc electron density maps for Arg residues contacting α- and γ-phosphate groups in a Y-like mode. Nucleotides and their analogs are shown as black sticks, interacting Arg residues as gray sticks. Mg^2+^ ions are shown as green spheres. Density map is colored according to the contouring level (1σ in pink , 2σ in blue, 3σ in green, 0.5σ in yellow) A. PDB 3ADD, chain B, Arg 116. Density contoured at 0.5σ and 1σ shown for phosphate chain and Arg residue. B. PDB 4TSF, Arg E356. Electron density shown for phosphate chain is contoured at 1σ and at 1σ and 2σ for Arg residue. C. PDB 4CEI B 283. Electron density shown for phosphate chain is contoured at 1σ and at 1σ, 2σ, 3σ for Arg residue. D. PDB 3LVD, Arg E469. Electron density shown for phosphate chain is contoured at 2σ and at 1σ and 2σ for Arg residue. E. PDB 3FFA, Arg A178. Density shown as in (B) F. PDB 3BB1, Arg G133. Density shown as in (B)

**Table SF1.**
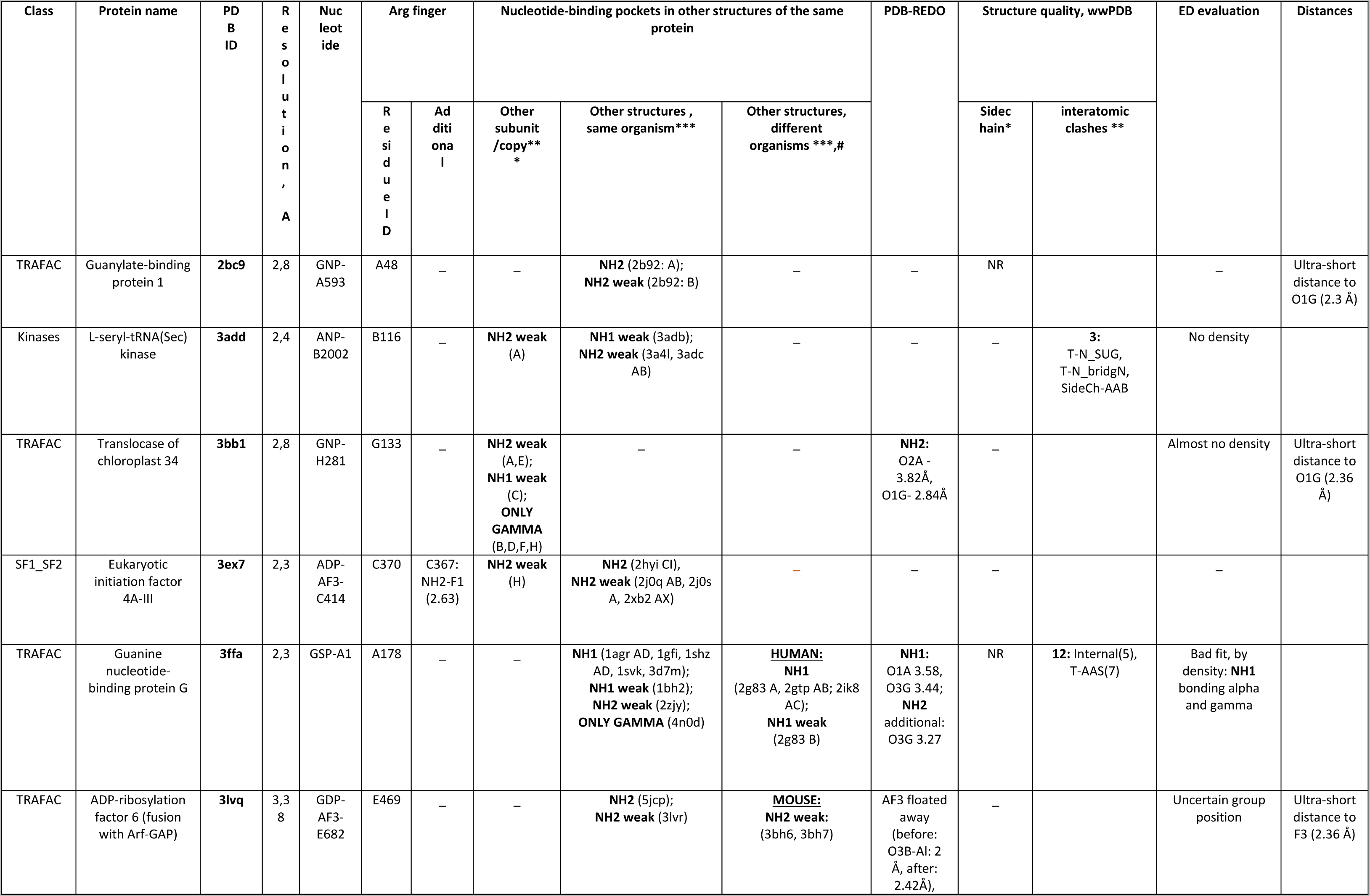
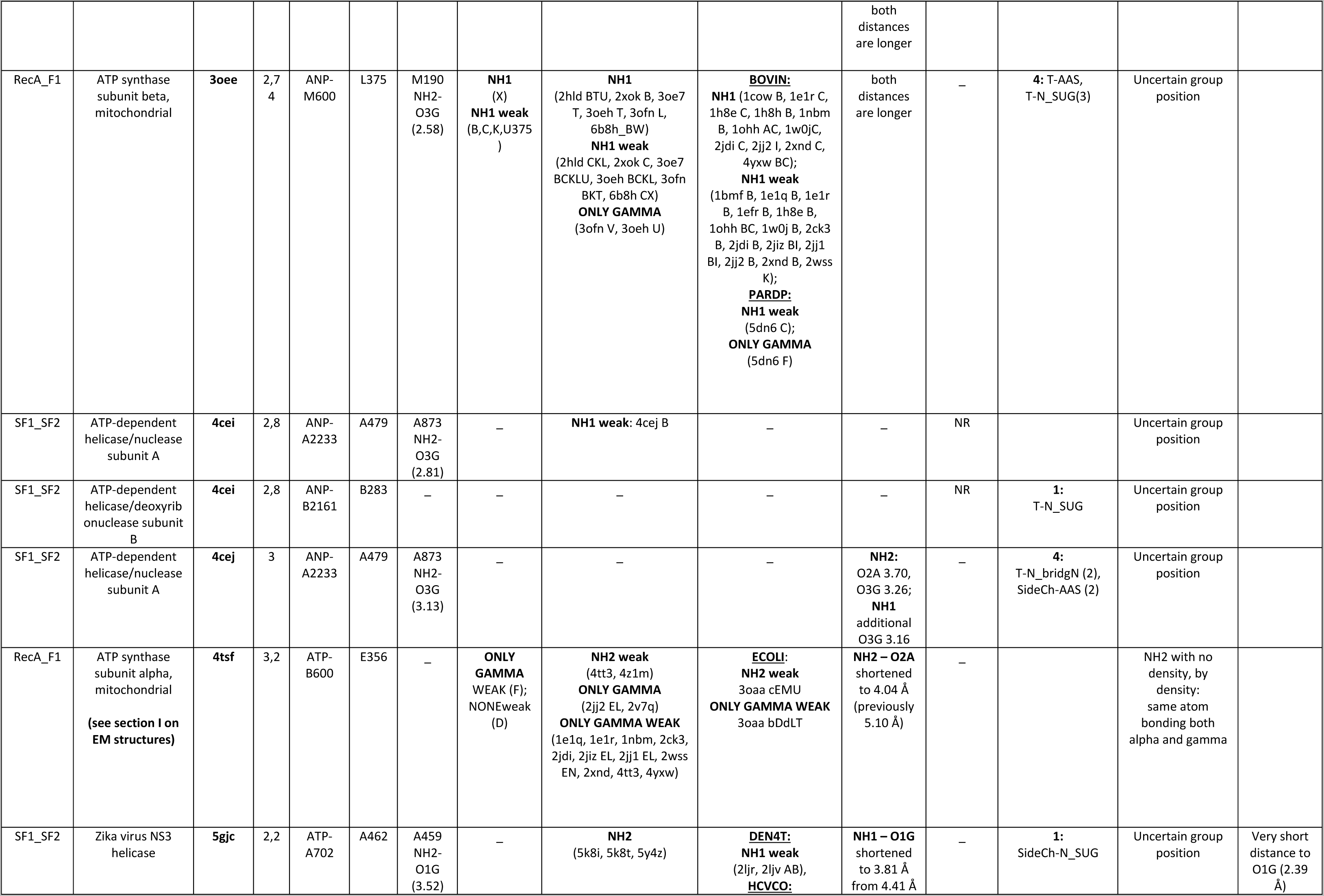
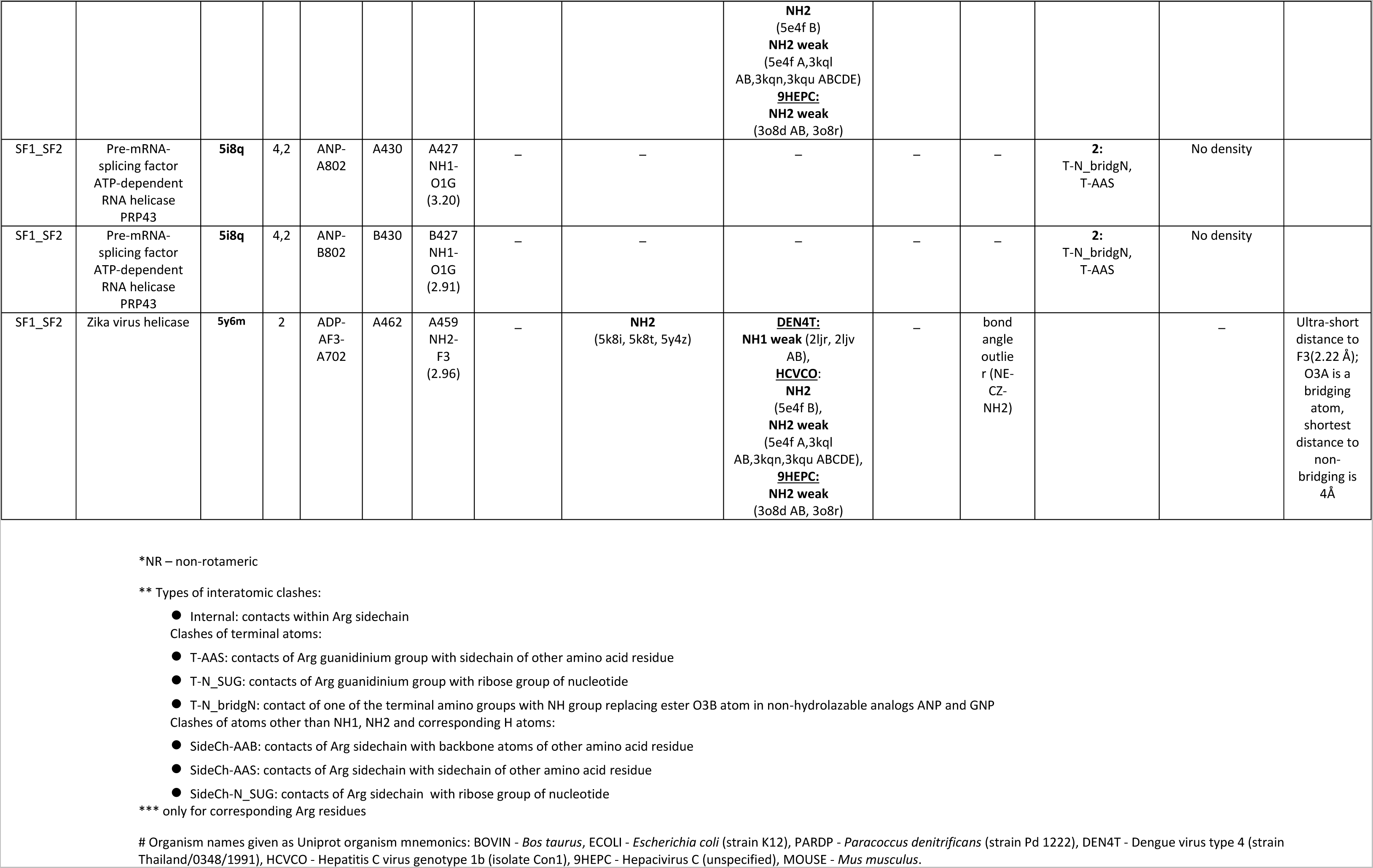
Arginine residues contacting phosphate chain in a Y-like mode.

## References

[1] J.E. Walker, M. Saraste, M.J. Runswick, N.J. Gay, Distantly related sequences in the alpha- and beta-subunits of ATP synthase, myosin, kinases and other ATP-requiring enzymes and a common nucleotide binding fold, EMBO J 1(8) (1982) 945–51.

[2] M. Saraste, P.R. Sibbald, A. Wittinghofer, The P-loop – a common motif in ATP- and GTP-binding proteins, Trends Biochem. Sci. 15(11) (1990) 430–4.

[3] A.E. Gorbalenya, E.V. Koonin, Helicases: amino acid sequence comparisons and structure-function relationships, Curr Opin Struc Biol 3(3) (1993) 419–429.

[4] C.A. Smith, I. Rayment, Active site comparisons highlight structural similarities between myosin and other P-loop proteins, Biophys J 70(4) (1996) 1590–602.

[5] A.F. Neuwald, L. Aravind, J.L. Spouge, E.V. Koonin, AAA+: A class of chaperone-like ATPases associated with the assembly, operation, and disassembly of protein complexes, Genome Res 9(1) (1999) 27–43.

[6] E. Muneyuki, H. Noji, T. Amano, T. Masaike, M. Yoshida, F(0)F(1)-ATP synthase: general structural features of ’ATP-engine’ and a problem on free energy transduction, Biochim Biophys Acta 1458(2-3) (2000) 467–81.

[7] D.D. Leipe, Y.I. Wolf, E.V. Koonin, L. Aravind, Classification and evolution of P-loop GTPases and related ATPases, J Mol Biol 317(1) (2002) 41–72.

[8] D.D. Leipe, E.V. Koonin, L. Aravind, Evolution and classification of P-loop kinases and related proteins, J Mol Biol 333(4) (2003) 781–815.

[9] V. Anantharaman, L. Aravind, E.V. Koonin, Emergence of diverse biochemical activities in evolutionarily conserved structural scaffolds of proteins, Curr. Opin. Chem. Biol. 7(1) (2003) 12–20.

[10] L.M. Iyer, D.D. Leipe, E.V. Koonin, L. Aravind, Evolutionary history and higher order classification of AAA+ ATPases, J. Struct. Biol. 146(1-2) (2004) 11–31.

[11] L.M. Iyer, K.S. Makarova, E.V. Koonin, L. Aravind, Comparative genomics of the FtsK-HerA superfamily of pumping ATPases: implications for the origins of chromosome segregation, cell division and viral capsid packaging, Nucleic Acids Res 32(17) (2004) 5260–79.

[12] A. Wittinghofer, I.R. Vetter, Structure-function relationships of the G domain, a canonical switch motif, Annu. Rev. Biochem. 80 (2011) 943–971.

[13] A.M. Burroughs, L. Aravind, The Origin and Evolution of Release Factors: Implications for Translation Termination, Ribosome Rescue, and Quality Control Pathways, Int J Mol Sci 20(8) (2019).

[14] L.M. Longo, J. Jablonska, P. Vyas, M. Kanade, R. Kolodny, N. Ben-Tal, D.S. Tawfik, On the emergence of P-Loop NTPase and Rossmann enzymes from a Beta-Alpha-Beta ancestral fragment, Elife 9 (2020).

[15] A. Krishnan, A.M. Burroughs, L.M. Iyer, L. Aravind, Comprehensive classification of ABC ATPases and their functional radiation in nucleoprotein dynamics and biological conflict systems, Nucleic Acids Res 48(18) (2020) 10045–10075.

[16] R.D. Schaeffer, Y. Liao, H. Cheng, N.V. Grishin, ECOD: new developments in the evolutionary classification of domains, Nucleic Acids Res 45(D1) (2017) D296–D302.

[17] R.D. Finn, P. Coggill, R.Y. Eberhardt, S.R. Eddy, J. Mistry, A.L. Mitchell, S.C. Potter, M. Punta, M. Qureshi, A. Sangrador-Vegas, G.A. Salazar, J. Tate, A. Bateman, The Pfam protein families database: towards a more sustainable future, Nucleic Acids Res. 44(D1) (2016) D279–D285.

[18] M.I. Kozlova, D.N. Shalaeva, D.V. Dibrova, A.Y. Mulkidjanian, Common mechanism of activated catalysis in P-loop fold nucleoside triphosphatases - united in diversity, BioRxiv (2022).

[19] A.N. Lupas, C.P. Ponting, R.B. Russell, On the evolution of protein folds: are similar motifs in different protein folds the result of convergence, insertion, or relics of an ancient peptide world?, J Struct Biol 134(2-3) (2001) 191–203.

[20] C.P. Ponting, R.R. Russell, The natural history of protein domains, Annual review of biophysics and biomolecular structure 31 (2002) 45–71.

[21] J. Söding, A.N. Lupas, More than the sum of their parts: on the evolution of proteins from peptides, BioEssays 25(9) (2003) 837–46.

[22] J.A. Ranea, A. Sillero, J.M. Thornton, C.A. Orengo, Protein superfamily evolution and the last universal common ancestor (LUCA), J. Mol. Evol. 63(4) (2006) 513–25.

[23] V. Alva, J. Soding, A.N. Lupas, A vocabulary of ancient peptides at the origin of folded proteins, eLife 4 (2015) e09410.

[24] M. Kanade, S. Chakraborty, S.S. Shelke, P. Gayathri, A Distinct Motif in a Prokaryotic Small Ras-Like GTPase Highlights Unifying Features of Walker B Motifs in P-Loop NTPases, J Mol Biol 432(20) (2020) 5544–5564.

[25] G.M. Blackburn, J. Cherfils, G.P. Moss, N.G.J. Richards, J.P. Waltho, N.H. Williams, A. Wittinghofer, How to name atoms in phosphates, polyphosphates, their derivatives and mimics, and transition state analogues for enzyme-catalysed phosphoryl transfer reactions (IUPAC Recommendations 2016), Pure Appl. Chem. 89(5) (2017) 653–675.

[26] X. Zhou, W. Ren, S.R. Bharath, X. Tang, Y. He, C. Chen, Z. Liu, D. Li, H. Song, Structural and Functional Insights into the Unwinding Mechanism of Bacteroides sp Pif1, Cell Rep 14(8) (2016) 2030–9.

[27] A. Scrima, A. Wittinghofer, Dimerisation-dependent GTPase reaction of MnmE: how potassium acts as GTPase-activating element, EMBO J 25(12) (2006) 2940–51.

[28] D.E. Coleman, A.M. Berghuis, E. Lee, M.E. Linder, A.G. Gilman, S.R. Sprang, Structures of active conformations of Gi alpha 1 and the mechanism of GTP hydrolysis, Science 265(5177) (1994) 1405–12.

[29] J. Sondek, D.G. Lambright, J.P. Noel, H.E. Hamm, P.B. Sigler, GTPase mechanism of Gproteins from the 1.7-A crystal structure of transducin alpha-GDP-AIF-4, Nature 372(6503) (1994) 276–9.

[30] K. Scheffzek, M.R. Ahmadian, W. Kabsch, L. Wiesmuller, A. Lautwein, F. Schmitz, A. Wittinghofer, The Ras-RasGAP complex: structural basis for GTPase activation and its loss in oncogenic Ras mutants, Science 277(5324) (1997) 333–8.

[31] K. Scheffzek, M.R. Ahmadian, A. Wittinghofer, GTPase-activating proteins: helping hands to complement an active site, Trends Biochem Sci 23(7) (1998) 257–62.

[32] T. Ogura, S.W. Whiteheart, A.J. Wilkinson, Conserved arginine residues implicated in ATP hydrolysis, nucleotide-sensing, and inter-subunit interactions in AAA and AAA+ ATPases, J. Structural Biology 146(1-2) (2004) 106–12.

[33] M.-R. Ash, M.J. Maher, J.M. Guss, M. Jormakka, The cation-dependent G-proteins: in a class of their own, FEBS Lett. 586(16) (2012) 2218–24.

[34] P. Wendler, S. Ciniawsky, M. Kock, S. Kube, Structure and function of the AAA+ nucleotide binding pocket, Biochim. Biophys. Acta 1823(1) (2012) 2–14.

[35] Y. Jin, R.W. Molt, Jr., G.M. Blackburn, Metal fluorides: Tools for structural and computational analysis of phosphoryl transfer enzymes, Top. Curr. Chem. (Cham) 375(2) (2017) 36.

[36] R. Gasper, F. Wittinghofer, The Ras switch in structural and historical perspective, Biol Chem 401(1) (2019) 143–163.

[37] I.R. Vetter, A. Wittinghofer, Signal transduction - The guanine nucleotide-binding switch in three dimensions, Science 294(5545) (2001) 1299–1304.

[38] K. Nam, J. Pu, M. Karplus, Trapping the ATP binding state leads to a detailed understanding of the F1-ATPase mechanism, Proc Natl Acad Sci U S A 111(50) (2014) 17851–6.

[39] P. Llinas, T. Isabet, L. Song, V. Ropars, B. Zong, H. Benisty, S. Sirigu, C. Morris, C. Kikuti, D. Safer, H.L. Sweeney, A. Houdusse, How actin initiates the motor activity of Myosin, Dev Cell 33(4) (2015) 401–12.

[40] A. Wittinghofer, Signaling mechanistics: aluminum fluoride for molecule of the year, Curr. Biol. 7(11) (1997) R682–R685.

[41] R.I. Menz, J.E. Walker, A.G. Leslie, Structure of bovine mitochondrial F_1_-ATPase with nucleotide bound to all three catalytic sites: implications for the mechanism of rotary catalysis, Cell 106(3) (2001) 331–41.

[42] D.L. Graham, P.N. Lowe, G.W. Grime, M. Marsh, K. Rittinger, S.J. Smerdon, S.J. Gamblin, J.F. Eccleston, MgF(3)(-) as a transition state analog of phosphoryl transfer, Chem Biol 9(3) (2002) 375–81.

[43] D.R. Davies, W.G. Hol, The power of vanadate in crystallographic investigations of phosphoryl transfer enzymes, FEBS Lett. 577(3) (2004) 315–21.

[44] Y. Jin, N.G. Richards, J.P. Waltho, G.M. Blackburn, Metal fluorides as analogues for studies on phosphoryl transfer enzymes, Angew. Chem. Int. Ed. Engl. 56(15) (2017) 4110–4128.

[45] J.R. Knowles, Enzyme-catalyzed phosphoryl transfer reactions, Annu Rev Biochem 49 (1980) 877–919.

[46] F.H. Westheimer, Why Nature Chose Phosphates, Science 235(4793) (1987) 1173–1178.

[47] Z.A. Shabarova, A.A. Bogdanov, Advanced Organic Chemistry of Nucleic Acids, VCH, Weinheim, 1994.

[48] M.W. Bowler, M.J. GCliff, J.P. Waltho, G.M. Blackburn, Why did Nature select phosphate for its dominant roles in biology?, New Journal of Chemistry 34 (2010) 784–794.

[49] J.K. Lassila, J.G. Zalatan, D. Herschlag, Biological phosphoryl-transfer reactions: understanding mechanism and catalysis, Annu Rev Biochem 80 (2011) 669–702.

[50] T. Higashijima, K.M. Ferguson, P.C. Sternweis, E.M. Ross, M.D. Smigel, A.G. Gilman, The effect of activating ligands on the intrinsic fluorescence of guanine nucleotide-binding regulatory proteins, J Biol Chem 262(2) (1987) 752–6.

[51] M. Chabre, Aluminofluoride and beryllofluoride complexes: new phosphate analogs in enzymology, Trends in biochemical sciences 15(1) (1990) 6–10.

[52] B. Antonny, J. Bigay, M. Chabre, A novel magnesium-dependent mechanism for the activation of transducin by fluoride, FEBS Lett 268(1) (1990) 277–80.

[53] B. Antonny, M. Sukumar, J. Bigay, M. Chabre, T. Higashijima, The mechanism of aluminum-independent G-protein activation by fluoride and magnesium. 31P NMR spectroscopy and fluorescence kinetic studies, J Biol Chem 268(4) (1993) 2393–402.

[54] I. Schlichting, J. Reinstein, pH influences fluoride coordination number of the AlFx phosphoryl transfer transition state analog, Nat Struct Biol 6(8) (1999) 721–3.

[55] D.L. Graham, J.F. Eccleston, C.W. Chung, P.N. Lowe, Magnesium fluoride-dependent binding of small G proteins to their GTPase-activating proteins, Biochemistry 38(45) (1999) 14981–7.

[56] M. Chaney, R. Grande, S.R. Wigneshweraraj, W. Cannon, P. Casaz, M.T. Gallegos, J. Schumacher, S. Jones, S. Elderkin, A.E. Dago, E. Morett, M. Buck, Binding of transcriptional activators to sigma 54 in the presence of the transition state analog ADP-aluminum fluoride: insights into activator mechanochemical action, Genes Dev 15(17) (2001) 2282–94.

[57] L. Gremer, B. Gilsbach, M.R. Ahmadian, A. Wittinghofer, Fluoride complexes of oncogenic Ras mutants to study the Ras-RasGap interaction, Biol. Chem. 389(9) (2008) 1163–71.

[58] N.J. Baxter, G.M. Blackburn, J.P. Marston, A.M. Hounslow, M.J. Cliff, W. Bermel, N.H. Williams, F. Hollfelder, D.E. Wemmer, J.P. Waltho, Anionic charge is prioritized over geometry in aluminum and magnesium fluoride transition state analogs of phosphoryl transfer enzymes, J Am Chem Soc 130(12) (2008) 3952–8.

[59] N. Zhang, M. Buck, Formation of MgF3 (-)-dependent complexes between an AAA(+) ATPase and sigma(54.), FEBS Open Bio 2 (2012) 89–92.

[60] T.M. Glennon, J. Villa, A. Warshel, How does GAP catalyze the GTPase reaction of Ras?: A computer simulation study, Biochemistry 39(32) (2000) 9641–9651.

[61] B.R. Prasad, N.V. Plotnikov, J. Lameira, A. Warshel, Quantitative exploration of the molecular origin of the activation of GTPase, Proc. Natl. Acad. Sci. USA 110(51) (2013) 20509–20514.

[62] S.C. Kamerlin, P.K. Sharma, R.B. Prasad, A. Warshel, Why nature really chose phosphate, Q. Rev. Biophys. 46(1) (2013) 1–132.

[63] C. Kotting, A. Kallenbach, Y. Suveyzdis, A. Wittinghofer, K. Gerwert, The GAP arginine finger movement into the catalytic site of Ras increases the activation entropy, Proc. Natl. Acad. Sci. USA 105(17) (2008) 6260–5.

[64] Y. Jin, R.W. Molt, Jr., J.P. Waltho, N.G. Richards, G.M. Blackburn, ^19^F NMR and DFT analysis reveal structural and electronic transition state features for RhoA-catalyzed GTP hydrolysis, Angew. Chem. Int. Ed. Engl. 55(10) (2016) 3318–22.

[65] R.W. Molt, Jr., E. Pellegrini, Y. Jin, A GAP-GTPase-GDP-Pi Intermediate Crystal Structure Analyzed by DFT Shows GTP Hydrolysis Involves Serial Proton Transfers, Chemistry 25(36) (2019) 8484–8488.

[66] K.A. Maegley, S.J. Admiraal, D. Herschlag, Ras-catalyzed hydrolysis of GTP: a new perspective from model studies, Proc. Natl. Acad. Sci. USA 93(16) (1996) 8160–6.

[67] T. Rudack, F. Xia, J. Schlitter, C. Kotting, K. Gerwert, Ras and GTPase-activating protein (GAP) drive GTP into a precatalytic state as revealed by combining FTIR and biomolecular simulations, Proc. Natl. Acad. Sci. USA 109(38) (2012) 15295–300.

[68] D. Mann, C. Teuber, S.A. Tennigkeit, G. Schroter, K. Gerwert, C. Kotting, Mechanism of the intrinsic arginine finger in heterotrimeric G proteins, Proc. Natl. Acad. Sci. USA 113(50) (2016) E8041–E8050.

[69] K. Gerwert, D. Mann, C. Kotting, Common mechanisms of catalysis in small and heterotrimeric GTPases and their respective GAPs, Biol. Chem. 398(5-6) (2017) 523–533.

[70] D.N. Shalaeva, D.A. Cherepanov, M.Y. Galperin, A.V. Golovin, A.Y. Mulkidjanian, Evolution of cation binding in the active sites of P-loop nucleoside triphosphatases in relation to the basic catalytic mechanism, Elife 7 (2018).

[71] K. Yamanaka, J. Hwang, M. Inouye, Characterization of GTPase activity of TrmE, a member of a novel GTPase superfamily, from *Thermotoga maritima*, J. Bacteriol. 182(24) (2000) 7078–82.

[72] S. Meyer, S. Bohme, A. Kruger, H.-J. Steinhoff, J.P. Klare, A. Wittinghofer, Kissing G domains of MnmE monitored by X-ray crystallography and pulse electron paramagnetic resonance spectroscopy, PLoS Biol. 7(10) (2009) e1000212.

[73] S. Bohme, S. Meyer, A. Kruger, H.J. Steinhoff, A. Wittinghofer, J.P. Klare, Stabilization of G domain conformations in the tRNA-modifying MnmE-GidA complex observed with double electron electron resonance spectroscopy, J. Biol. Chem. 285(22) (2010) 16991–7000.

[74] B. Anand, P. Surana, B. Prakash, Deciphering the catalytic machinery in 30S ribosome assembly GTPase YqeH, PLoS One 5(4) (2010) e9944.

[75] J.A. Ballestros, H. Weinstein, Integrated methods for the construction of three-dimensional models and computational probing of structure-function relations in G protein-coupled receptors, Methods Neurosci. 25 (1995) 366–428.

[76] M.V. Milburn, L. Tong, A.M. deVos, A. Brunger, Z. Yamaizumi, S. Nishimura, S.H. Kim, Molecular switch for signal transduction: structural differences between active and inactive forms of protooncogenic ras proteins, Science 247(4945) (1990) 939–45.

[77] M. Frech, J. John, V. Pizon, P. Chardin, A. Tavitian, R. Clark, F. McCormick, A. Wittinghofer, Inhibition of GTPase activating protein stimulation of Ras-p21 GTPase by the Krev-1 gene product, Science 249(4965) (1990) 169–71.

[78] H.M. Berman, J. Westbrook, Z. Feng, G. Gilliland, T.N. Bhat, H. Weissig, I.N. Shindyalov, P.E. Bourne, The Protein Data Bank, Nucleic Acids Res 28(1) (2000) 235–42.

[79] S.K. Burley, C. Bhikadiya, C. Bi, S. Bittrich, L. Chen, G.V. Crichlow, C.H. Christie, K. Dalenberg, L. Di Costanzo, J.M. Duarte, S. Dutta, Z. Feng, S. Ganesan, D.S. Goodsell, S. Ghosh, R.K. Green, V. Guranovic, D. Guzenko, B.P. Hudson, C.L. Lawson, Y. Liang, R. Lowe, H. Namkoong, E. Peisach, I. Persikova, C. Randle, A. Rose, Y. Rose, A. Sali, J. Segura, M. Sekharan, C. Shao, Y.P. Tao, M. Voigt, J.D. Westbrook, J.Y. Young, C. Zardecki, M. Zhuravleva, RCSB Protein Data Bank: powerful new tools for exploring 3D structures of biological macromolecules for basic and applied research and education in fundamental biology, biomedicine, biotechnology, bioengineering and energy sciences, Nucleic Acids Res 49(D1) (2021) D437–D451.

[80] M. Blum, H.Y. Chang, S. Chuguransky, T. Grego, S. Kandasaamy, A. Mitchell, G. Nuka, T. Paysan-Lafosse, M. Qureshi, S. Raj, L. Richardson, G.A. Salazar, L. Williams, P. Bork, A. Bridge, J. Gough, D.H. Haft, I. Letunic, A. Marchler-Bauer, H. Mi, D.A. Natale, M. Necci, C.A. Orengo, A.P. Pandurangan, C. Rivoire, C.J.A. Sigrist, I. Sillitoe, N. Thanki, P.D. Thomas, S.C.E. Tosatto, C.H. Wu, A. Bateman, R.D. Finn, The InterPro protein families and domains database: 20 years on, Nucleic Acids Res 49(D1) (2021) D344–D354.

[81] E. Martz, Help, Index & Glossary for Protein Explorer, http://www.umass.edu/microbio/chime/pe_beta/pe/protexpl/igloss.htm?q=microbio/chime/explorer/igloss.htm, 2001.

[82] V.S. Minkov, V.V. Ghazaryan, E.V. Boldyreva, A.M. Petrosyan, Unusual hydrogen bonding in L-cysteine hydrogen fluoride, Acta Crystallogr C Struct Chem 71(Pt 8) (2015) 733–41.

[83] G.A. Jeffrey, An introduction to hydrogen bonding, Oxford University Press1997.

[84] R.C. Yu, P.I. Hanson, R. Jahn, A.T. Brunger, Structure of the ATP-dependent oligomerization domain of N-ethylmaleimide sensitive factor complexed with ATP, Nat Struct Biol 5(9) (1998) 803–11.

[85] W. Wu, K.T. Park, T. Holyoak, J. Lutkenhaus, Determination of the structure of the MinD-ATP complex reveals the orientation of MinD on the membrane and the relative location of the binding sites for MinE and MinC, Mol Microbiol 79(6) (2011) 1515–28.

[86] T.A. Leonard, P.J. Butler, J. Lowe, Bacterial chromosome segregation: structure and DNA binding of the Soj dimer--a conserved biological switch, EMBO J 24(2) (2005) 270–82.

[87] X. Yang, C. Chen, H. Tian, H. Chi, Z. Mu, T. Zhang, K. Yang, Q. Zhao, X. Liu, Z. Wang, X. Ji, H. Yang, Mechanism of ATP hydrolysis by the Zika virus helicase, Faseb J 32(10) (2018) 5250–5257.

[88] C.A. Orengo, J.M. Thornton, Protein families and their evolution-a structural perspective, Annu. Rev. Biochem. 74 (2005) 867–900.

[89] M. Gu, C.M. Rice, The Spring alpha-Helix Coordinates Multiple Modes of HCV (Hepatitis C Virus) NS3 Helicase Action, J Biol Chem 291(28) (2016) 14499–509.

[90] E. Uchikawa, M. Lethier, H. Malet, J. Brunel, D. Gerlier, S. Cusack, Structural Analysis of dsRNA Binding to Anti-viral Pattern Recognition Receptors LGP2 and MDA5, Mol Cell 62(4) (2016) 586–602.

[91] M. Soundararajan, F.S. Willard, A.J. Kimple, A.P. Turnbull, L.J. Ball, G.A. Schoch, C. Gileadi, O.Y. Fedorov, E.F. Dowler, V.A. Higman, S.Q. Hutsell, M. Sundstrom, D.A. Doyle, D.P. Siderovski, Structural diversity in the RGS domain and its interaction with heterotrimeric G protein alpha-subunits, Proc Natl Acad Sci U S A 105(17) (2008) 6457–62.

[92] D.N. Shalaeva, D.A. Cherepanov, M.Y. Galperin, A.Y. Mulkidjanian, Comparative analysis of active sites in P-loop nucleoside triphosphatases suggests an ancestral activation mechanism, BioRxiv 439992 [preprint] (2018).

[93] M.L. Oldham, J. Chen, Snapshots of the maltose transporter during ATP hydrolysis, Proc Natl Acad Sci U S A 108(37) (2011) 15152–6.

[94] F. Yi, R. Kong, J. Ren, L. Zhu, J. Lou, J.Y. Wu, W. Feng, Noncanonical Myo9b-RhoGAP Accelerates RhoA GTP Hydrolysis by a Dual-Arginine-Finger Mechanism, J Mol Biol 428(15) (2016) 3043–57.

[95] V.G. Taylor, P.A. Bommarito, J.J. Tesmer, Structure of the Regulator of G Protein Signaling 8 (RGS8)-Galphaq Complex: MOLECULAR BASIS FOR Galpha SELECTIVITY, J Biol Chem 291(10) (2016) 5138–45.

[96] J. Abe, T.B. Hiyama, A. Mukaiyama, S. Son, T. Mori, S. Saito, M. Osako, J. Wolanin, E. Yamashita, T. Kondo, S. Akiyama, Circadian rhythms. Atomic-scale origins of slowness in the cyanobacterial circadian clock, Science 349(6245) (2015) 312–6.

[97] N.L. Jean, T.J. Rutherford, J. Lowe, FtsK in motion reveals its mechanism for double-stranded DNA translocation, Proc Natl Acad Sci U S A 117(25) (2020) 14202–14208.

[98] D. Gai, R. Zhao, D. Li, C.V. Finkielstein, X.S. Chen, Mechanisms of conformational change for a replicative hexameric helicase of SV40 large tumor antigen, Cell 119(1) (2004) 47–60.

[99] A. Mateja, A. Szlachcic, M.E. Downing, M. Dobosz, M. Mariappan, R.S. Hegde, R.J. Keenan, The structural basis of tail-anchored membrane protein recognition by Get3, Nature 461(7262) (2009) 361–6.

[100] J.S. Chappie, S. Acharya, M. Leonard, S.L. Schmid, F. Dyda, G domain dimerization controls dynamin’s assembly-stimulated GTPase activity, Nature 465(7297) (2010) 435–40.

[101] X. Qian, Y. He, Y. Wu, Y. Luo, Asp302 determines potassium dependence of a RadA recombinase from *Methanococcus voltae*, J. Mol. Biol. 360(3) (2006) 537–47.

[102] D.V. Dibrova, M.Y. Galperin, E.V. Koonin, A.Y. Mulkidjanian, Ancient systems of sodium/potassium homeostasis as predecessors of membrane bioenergetics, Biochemistry. Biokhimiia 80(5) (2015) 495–516.

[103] D.C. Rees, E. Johnson, O. Lewinson, ABC transporters: the power to change, Nat Rev Mol Cell Biol 10(3) (2009) 218–27.

[104] M. Dean, A. Rzhetsky, R. Allikmets, The human ATP-binding cassette (ABC) transporter superfamily, Genome Res 11(7) (2001) 1156–66.

[105] I.D. Kerr, Sequence analysis of twin ATP binding cassette proteins involved in translational control, antibiotic resistance, and ribonuclease L inhibition, Biochem Biophys Res Commun 315(1) (2004) 166–73.

[106] K.P. Hopfner, A. Karcher, D.S. Shin, L. Craig, L.M. Arthur, J.P. Carney, J.A. Tainer, Structural biology of Rad50 ATPase: ATP-driven conformational control in DNA double-strand break repair and the ABC-ATPase superfamily, Cell 101(7) (2000) 789–800.

[107] J.A. Eisen, A phylogenomic study of the MutS family of proteins, Nucleic Acids Res 26(18) (1998) 4291–300.

[108] A. Decottignies, A. Goffeau, Complete inventory of the yeast ABC proteins, Nat Genet 15(2) (1997) 137–45.

[109] K.A. Schug, W. Lindner, Noncovalent binding between guanidinium and anionic groups: focus on biological- and synthetic-based arginine/guanidinium interactions with phosph[on]ate and sulf[on]ate residues, Chem Rev 105(1) (2005) 67–114.

[110] B.J. Calnan, B. Tidor, S. Biancalana, D. Hudson, A.D. Frankel, Arginine-mediated RNA recognition: the arginine fork, Science 252(5009) (1991) 1167–71.

[111] A.V. Afonin, I.V. Sterkhova, A.V. Vashchenko, M.V. Sigalov, Estimating the energy of intramolecular bifurcated (three-centered) hydrogen bond by X-ray, IR and 1H NMR spectroscopy, and QTAIM calculations, J Mol Struct 1163 (2018) 185–196.

[112] D. Lacabanne, T. Wiegand, N. Wili, M.I. Kozlova, R. Cadalbert, D. Klose, A.Y. Mulkidjanian, B.H. Meier, A. Bockmann, ATP Analogues for Structural Investigations: Case Studies of a DnaB Helicase and an ABC Transporter, Molecules 25(22) (2020).

[113] A.A. Malaer, N. Wili, L.A. Volker, M.I. Kozlova, R. Cadalbert, A. Dapp, M.E. Weber, J. Zehnder, G. Jeschke, H. Eckert, A. Bockmann, D. Klose, A.Y. Mulkidjanian, B.H. Meier, T. Wiegand, Spectroscopic glimpses of the transition state of ATP hydrolysis trapped in a bacterial DnaB helicase, Nat Commun 12(1) (2021) 5293.

[114] A.J. Fisher, C.A. Smith, J.B. Thoden, R. Smith, K. Sutoh, H.M. Holden, I. Rayment, X-ray structures of the myosin motor domain of Dictyostelium discoideum complexed with MgADP.BeFx and MgADP.AlF4, Biochemistry 34(28) (1995) 8960–72.

[115] M.A. Geeves, Review: The ATPase mechanism of myosin and actomyosin, Biopolymers 105(8) (2016) 483–91.

[116] R.A. Cross, Review: Mechanochemistry of the kinesin-1 ATPase, Biopolymers 105(8) (2016) 476–82.

[117] A.S. Woods, S. Ferre, Amazing stability of the arginine-phosphate electrostatic interaction, J Proteome Res 4(4) (2005) 1397–402.

[118] L. Shimoni, J.P. Glusker, Hydrogen bonding motifs of protein side chains: descriptions of binding of arginine and amide groups, Protein Sci 4(1) (1995) 65–74.

[119] C.A. Seipp, N.J. Williams, M.K. Kidder, R. Custelcean, CO2 Capture from Ambient Air by Crystallization with a Guanidine Sorbent, Angew Chem Int Ed Engl 56(4) (2017) 1042–1045.

[120] I.K. McDonald, J.M. Thornton, Satisfying hydrogen bonding potential in proteins, J Mol Biol 238(5) (1994) 777–93.

[121] B. van Beusekom, W.G. Touw, M. Tatineni, S. Somani, G. Rajagopal, J. Luo, G.L. Gilliland, A. Perrakis, R.P. Joosten, Homology-based hydrogen bond information improves crystallographic structures in the PDB, Protein Sci 27(3) (2018) 798–808.

[122] D. Sehnal, S. Bittrich, M. Deshpande, R. Svobodova, K. Berka, V. Bazgier, S. Velankar, S.K. Burley, J. Koca, A.S. Rose, Mol* Viewer: modern web app for 3D visualization and analysis of large biomolecular structures, Nucleic Acids Res 49(W1) (2021) W431–W437.

[123] W.L. DeLano, The PyMOL Molecular Graphics System, Version 1.7.2.1, Schrödinger, LLC., 2010.

## References

[1] D.D. Leipe, Y.I. Wolf, E.V. Koonin, L. Aravind, Classification and evolution of P-loop GTPases and related ATPases, J Mol Biol 317(1) (2002) 41–72.

[2] D.D. Leipe, E.V. Koonin, L. Aravind, Evolution and classification of P-loop kinases and related proteins, J Mol Biol 333(4) (2003) 781–815.

[3] L.M. Iyer, D.D. Leipe, E.V. Koonin, L. Aravind, Evolutionary history and higher order classification of AAA+ ATPases, J. Struct. Biol. 146(1-2) (2004) 11–31.

[4] L.M. Iyer, K.S. Makarova, E.V. Koonin, L. Aravind, Comparative genomics of the FtsK-HerA superfamily of pumping ATPases: implications for the origins of chromosome segregation, cell division and viral capsid packaging, Nucleic Acids Res 32(17) (2004) 5260–79.

[5] D.D. Leipe, E.V. Koonin, L. Aravind, STAND, a class of P-loop NTPases including animal and plant regulators of programmed cell death: multiple, complex domain architectures, unusual phyletic patterns, and evolution by horizontal gene transfer, J. Mol. Biol. 343(1) (2004) 1–28.

[6] D.N. Shalaeva, D.A. Cherepanov, M.Y. Galperin, A.V. Golovin, A.Y. Mulkidjanian, Evolution of cation binding in the active sites of P-loop nucleoside triphosphatases in relation to the basic catalytic mechanism, Elife 7 (2018).

## References

[1] J. Gu, L. Zhang, S. Zong, R. Guo, T. Liu, J. Yi, P. Wang, W. Zhuo, M. Yang, Cryo-EM structure of the mammalian ATP synthase tetramer bound with inhibitory protein IF1, Science 364(6445) (2019) 1068–1075.

[2] B.J. Murphy, N. Klusch, J. Langer, D.J. Mills, O. Yildiz, W. Kuhlbrandt, Rotary substates of mitochondrial ATP synthase reveal the basis of flexible F1-Fo coupling, Science 364(6446) (2019).

[3] N. Kapoor, S.T. Menon, R. Chauhan, P. Sachdev, T.P. Sakmar, Structural evidence for a sequential release mechanism for activation of heterotrimeric G proteins, J Mol Biol 393(4) (2009) 882–97.

[4] K. Scheffzek, M.R. Ahmadian, W. Kabsch, L. Wiesmuller, A. Lautwein, F. Schmitz, A. Wittinghofer, The Ras-RasGAP complex: structural basis for GTPase activation and its loss in oncogenic Ras mutants, Science 277(5324) (1997) 333–8.

[5] D.E. Coleman, A.M. Berghuis, E. Lee, M.E. Linder, A.G. Gilman, S.R. Sprang, Structures of active conformations of Gi alpha 1 and the mechanism of GTP hydrolysis, Science 265(5177) (1994) 1405–12.

[6] D.L. Graham, P.N. Lowe, G.W. Grime, M. Marsh, K. Rittinger, S.J. Smerdon, S.J. Gamblin, J.F. Eccleston, MgF(3)(-) as a transition state analog of phosphoryl transfer, Chem Biol 9(3) (2002) 375–81.

[7] K. Rittinger, P.A. Walker, J.F. Eccleston, S.J. Smerdon, S.J. Gamblin, Structure at 1.65 A of RhoA and its GTPase-activating protein in complex with a transition-state analogue, Nature 389(6652) (1997) 758–62.

[8] E. Amin, M. Jaiswal, U. Derewenda, K. Reis, K. Nouri, K.T. Koessmeier, P. Aspenstrom, A.V. Somlyo, R. Dvorsky, M.R. Ahmadian, Deciphering the Molecular and Functional Basis of RHOGAP Family Proteins: A SYSTEMATIC APPROACH TOWARD SELECTIVE INACTIVATION OF RHO FAMILY PROTEINS, J Biol Chem 291(39) (2016) 20353–71.

[9] H. Resat, T.P. Straatsma, D.A. Dixon, J.H. Miller, The arginine finger of RasGAP helps Gln-61 align the nucleophilic water in GAP-stimulated hydrolysis of GTP, Proc. Natl. Acad. Sci. USA 98(11) (2001) 6033–8.

[10] R.P. Joosten, J. Salzemann, V. Bloch, H. Stockinger, A.C. Berglund, C. Blanchet, E. Bongcam-Rudloff, C. Combet, A.L. Da Costa, G. Deleage, M. Diarena, R. Fabbretti, G. Fettahi, V. Flegel, A. Gisel, V. Kasam, T. Kervinen, E. Korpelainen, K. Mattila, M. Pagni, M. Reichstadt, V. Breton, I.J. Tickle, G. Vriend, PDB_REDO: automated re-refinement of X-ray structure models in the PDB, J Appl Crystallogr 42(Pt 3) (2009) 376–384.

